# IRON REGULATORY PROTEIN (IRP)-MEDIATED IRON HOMEOSTASIS IS CRITICAL FOR NEUTROPHIL DEVELOPMENT AND DIFFERENTIATION IN THE BONE MARROW

**DOI:** 10.1101/2022.03.16.483971

**Authors:** Michael Bonadonna, Sandro Altamura, Elisabeth Tybl, Gael Palais, Maria Qatato, Maria Polycarpou-Schwarz, Martin Schneider, Christina Kalk, Wibke Rüdiger, Alina Ertl, Natasha Anstee, Ruzhica Bogeska, Dominic Helm, Michael D. Milsom, Bruno Galy

## Abstract

Iron is mostly devoted to the hemoglobinization of erythrocytes for oxygen transport. Yet, emerging evidence points to a broader role for the metal in hematopoiesis, including the formation of the immune system. Iron availability in mammalian cells is controlled by iron-regulatory proteins (IRP)-1 and −2. We report that global disruption of both IRP1 and IRP2 in adult mice impairs neutrophil development and differentiation in the bone marrow, yielding immature neutrophils with abnormally high glycolytic and autophagic activity, resulting in neutropenia. IRPs promote neutrophil differentiation in a cell intrinsic manner by securing cellular iron supply together with transcriptional control of neutropoiesis to facilitate differentiation to fully mature neutrophils. Unlike neutrophils, monocyte count was not affected by IRP and iron deficiency, suggesting a lineage-specific effect of iron on myeloid output. This study unveils the previously unrecognized importance of IRPs and iron metabolism in the formation of a major branch of the innate immune system.

## INTRODUCTION

As an integral component of heme, iron must be supplied in adequate amounts to secure hemoglobin synthesis and prevent anemia. Anemia due to iron deficiency is highly prevalent (Pasricha et al., 2021). It is often due to insufficient dietary iron intake, or to inhibition of iron delivery into the plasma in conditions of inflammation (Ganz, 2019, Haschka et al., 2021). Iron overload resulting from e.g. ineffective erythropoiesis or mutations in iron homeostasis genes, is also deleterious, affecting organs such as the liver and heart, and promoting infections.

Body iron levels and distribution are controlled by the peptide hepcidin (or HAMP). Hepcidin reduces the release of iron into the plasma from iron exporting cells by binding to and inhibiting the iron exporter ferroportin (FPN, also known as SLC40A1). Its levels augment in response to immune activation or when body iron levels are high. Conversely, stimulation of red blood cell (RBC) production suppresses hepcidin via the erythroblast-derived hormone erythroferrone (ERFE, also FAM132B), allowing for iron export via FPN to match erythroid iron needs (Muckenthaler et al., 2017).

At the cellular level, iron homeostasis is orchestrated by the interaction of iron regulatory proteins (IRP)-1 and −2 (also known as *ACO1* and *IREB2*, respectively) with *cis*-regulatory iron responsive elements (IREs) present in the untranslated region (UTR) of mRNAs encoding iron metabolism molecules. When iron is scarce, IRPs bind to the 5’-UTR IRE of mRNAs encoding the ferritin-H and −L (FTH1 and FTL1, respectively) iron storage proteins, the iron exporter FPN, the erythroid aminolevulinate synthase (ALAS2), or the transcription hypoxia inducible factor 2 alpha (HIF2A, also EPAS1) and inhibit their translation. IRP binding to 3’-UTR IREs in the transferrin receptor 1 (TFRC) mRNA protects the transcript against the Regnase-1 (ZC3H12A) and Roquin-1 (RC3H1) nucleases (Yoshinaga et al., 2017, Corral et al., 2021), thereby promoting iron uptake. In iron and oxygen-replete cells, IRP2 is displaced from IRE RNA by F-box and leucine rich repeat protein 5 (FBXL5) and is targeted for degradation (Wang et al., 2020), whereas IRP1 assembles an iron-sulfur cluster (ISC) and functions as an aconitase (Rouault and Maio, 2017).

Studies in patients and animal models revealed that inadequate IRP activity is detrimental. Indeed, abnormal stimulation of IRP activity due to defects in ISC biogenesis impairs heme synthesis in erythroid cells (Ward and Cloonan, 2019), and aberrant gain of IRP2 function in FBXL5-null mice was found to be lethal (Moroishi et al., 2011). Conversely, constitutive, systemic deletion of IRP2 in mice results in microcytic anemia, diabetes, and neurological symptoms of varying severity depending on the mouse strain examined (Wilkinson and Pantopoulos, 2014, Santos et al., 2020). IRP1 deficiency results in transient polycythemia during early life, due to derepression of *Hif2a* translation in kidney cells and subsequent stimulation of erythropoietin (*Epo)*. Importantly, despite these phenotypic traits, mice lacking either of the two IRPs remain viable and fertile. This is in distinct contrast with the early embryonic lethality resulting from double IRP deficiency (Smith et al., 2006, Galy et al., 2008), and shows that while the IRP/IRE network is essential, the two IRPs can largely compensate for each other.

We set out to determine the functions of the IRP/IRE regulatory network during adult life. To circumvent the early lethality due to compound IRP deficiency, we took advantage of conditional *Irp* alleles to acutely disrupt both IRP1 and IRP2 in the entire body of adult mice. Our study reveals that IRPs are critically important for granulocyte development and differentiation in the bone marrow, unveiling a previously unrecognized role for iron metabolism and the IRP/IRE regulatory network in hematopoiesis and formation of the immune system.

## RESULTS

### Systemic acute loss of IRP function causes erythropenia and myelopenia

To study the impact of IRP deficiency during adult life, we generated mice that carry floxed *Aco1* and *Ireb2* alleles (Galy et al., 2005a) and express a tamoxifen-inducible CRE recombinase (CreER) under the control of the *Rosa26* promoter (Badea et al., 2003). The resulting mice are designated P1/2-KO; control littermates lacking CreER are designated P1/2-CTR. To control for potential effects of CRE (Hameyer et al., 2007, Higashi et al., 2009), we also analyzed animals carrying the CreER sequence alone (CreER) and their wild-type littermates (WT). Adult P1/2-KO mice treated with two non-consecutive low doses of tamoxifen displayed efficient recombination of the floxed *Aco1* and *Ireb2* alleles in most tissues analyzed except for brain (Figure 1A).

**Figure 1:**
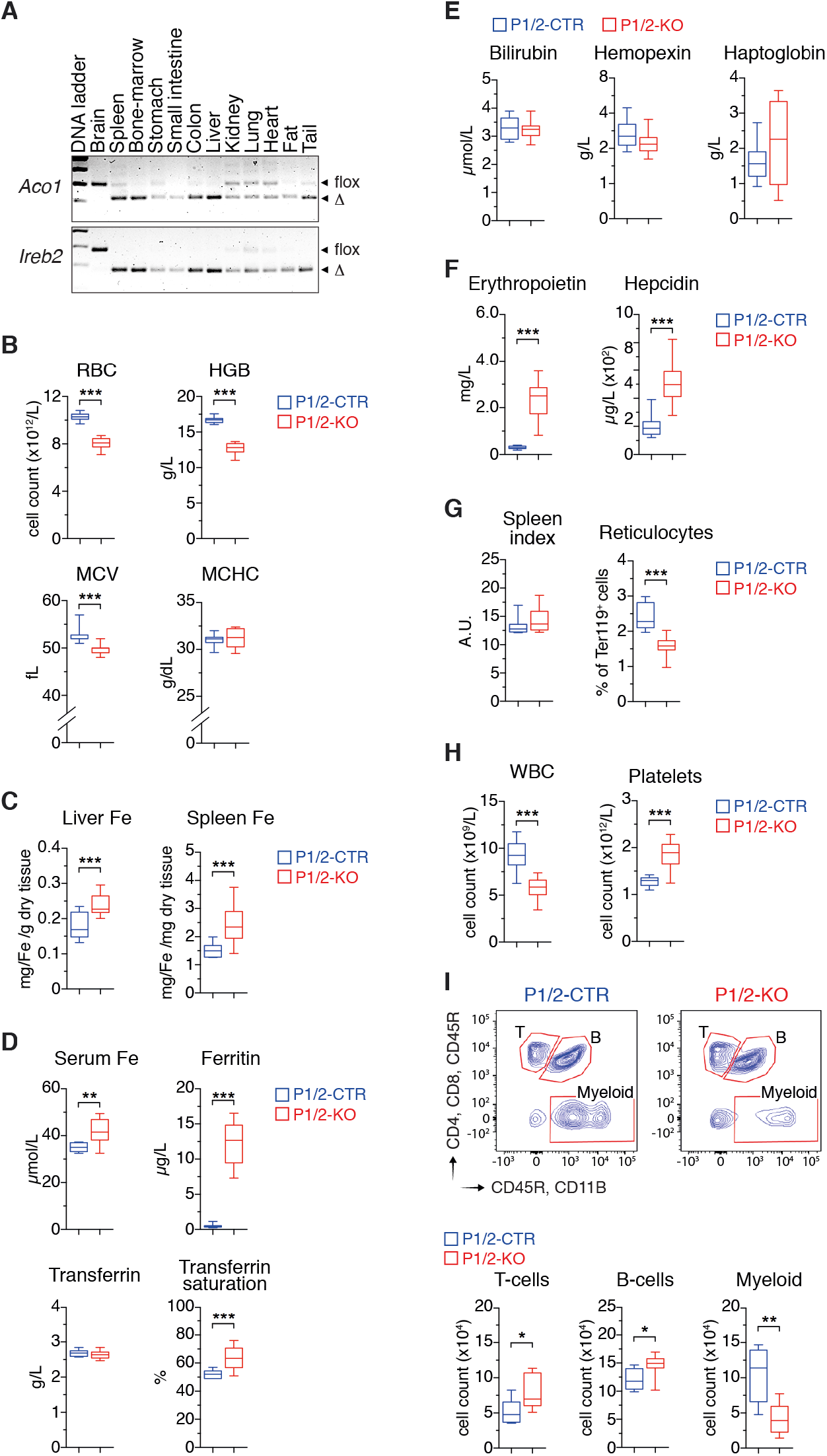
Acute IRP ablation during adulthood causes erythropenia and myelopenia. P1/2-CTR and P1/2-KO littermates received tamoxifen on day 1 and day 3 to induce IRP ablation. Mice were analyzed on day 10. (A) Genomic PCR analysis of the *Irp1* (*Aco1*) and *Irp2* (*Ireb2*) alleles in tissues from P1/2-KO mice. The floxed (flox) and truncated (Δ) alleles are indicated. (B) Red blood cell (RBC) parameters (HGB: hemoglobin; MCV: mean corpuscular volume; MCHC: mean corpuscular hemoglobin concentration). (C) Hepatic and splenic iron levels. (D) Serum iron parameters. (E) RBC decay markers. (F) Serum levels of erythropoietin and hepcidin. (G) Spleen index= √[(100Xspleen weight in mg/body weigh t in g)], and reticulocyte frequency in peripheral blood (PB). (H) White blood cell (WBC) and platelet counts in PB. (I) Flow cytometry (FCM) analysis of major WBC populations in PB. The gating strategy is shown on contour plots on the left. The histograms display cell counts for 3×10^5^ events recorded. B-I) The results are presented as box plots (minimum to maximum values). Sample size: P1/2-KO and P1/2-CTR, n=15-18 (B, F, H), n=7-12 (C-E, I). * p<0.05; ** p<0.01; *** p<0.001. *See also Figures S1 and S2.*

On day 10 after the commencement of tamoxifen treatment, P1/2-KO mice exhibited normal overall posture and activity pattern, and their body weight was unchanged. However, these mice display a substantial decrease in the number and size of red blood cells (RBCs), as well as low hemoglobin levels (Figure 1B). The anemia was accompanied by iron accumulation in the liver and spleen, elevated plasma iron and ferritin levels, and high transferrin saturation (Figure 1C,D). Despite high iron levels, IRP-deficient mice were anemic, suggesting that the anemia may be due to hemolysis and/or deficient RBC production. However, hemolysis is unlikely because serum bilirubin, haptoglobin, and hemopexin levels are unchanged (Figure 1E). Of note, serum EPO concentration was markedly increased in P1/2-KO animals (Figure 1F). High EPO levels are predicted to stimulate erythropoiesis and suppress hepcidin via ERFE. However, we observed a decrease in blood reticulocytes, and no sign of extramedullary erythropoiesis (Figure 1G). Moreover, ERFE remained below the level of detection (not shown) and hepcidin concentration was increased (Figure 1F). Together, these results indicated that loss of IRP function suppresses erythropoiesis, thereby decreasing iron consumption and leading to iron accumulation in the body.

Surprisingly, IRP ablation not only altered RBC numbers but also resulted in a substantial reduction in white blood cells (WBC, Figure 1H), a feature not observed in mice with single IRP1 or IRP2 ablation (Figure S1); platelets counts were increased (Figure 1H). To better define what type of WBC is affected by IRP-deficiency, we analyzed the major leukocyte populations in peripheral blood (PB) by flow cytometry (FCM). P1/2-KO animals displayed a 60% decrease in the number of myeloid cells, whereas B- and T- cells were slightly increased (Figure 1I). Importantly, the alterations in erythrocyte and leukocyte parameters were not consequences of CRE activation, as CreER animals subjected to the same regimen were largely asymptomatic (Figure S2A-F).

Collectively, these results revealed that, in addition to their critical role in erythropoiesis, IRPs are essential for maintaining physiological levels of myeloid cells during adult life.

### Loss of IRP function impairs the production of erythrocytes and granulocytes

We speculated that the reduction in erythroid and myeloid cells in P1/2-KO animals might be due to impaired hematopoiesis in the bone marrow (BM). Consistent with this hypothesis, we observed a 50 % decrease in BM cellularity in P1/2-KO versus P1/2-CTR animals (Figure 2A). Hematopoiesis is typically depicted as a hierarchical process, in which self-renewing multipotent long-term (LT) hematopoietic stem cells (HSCs) give rise to short-term (ST)-HSCs that progress to lineage-committed progenitors with increasingly limited differentiation and self-renewal capacity, and finally to terminally-differentiated hematopoietic cells (Figure 2B). P1/2-KO mice exhibited a clear expansion of both LT- and ST-HSCs, accompanied by an increase in all sub-types of multipotent progenitors (MPPs, Figure 2C). The amount of common myeloid (CMP), megakaryocyte/erythroid (MEP), granulocyte/monocyte (GMP), and common lymphoid progenitors (CLP) was also elevated (Figure 2D).

**Figure 2:**
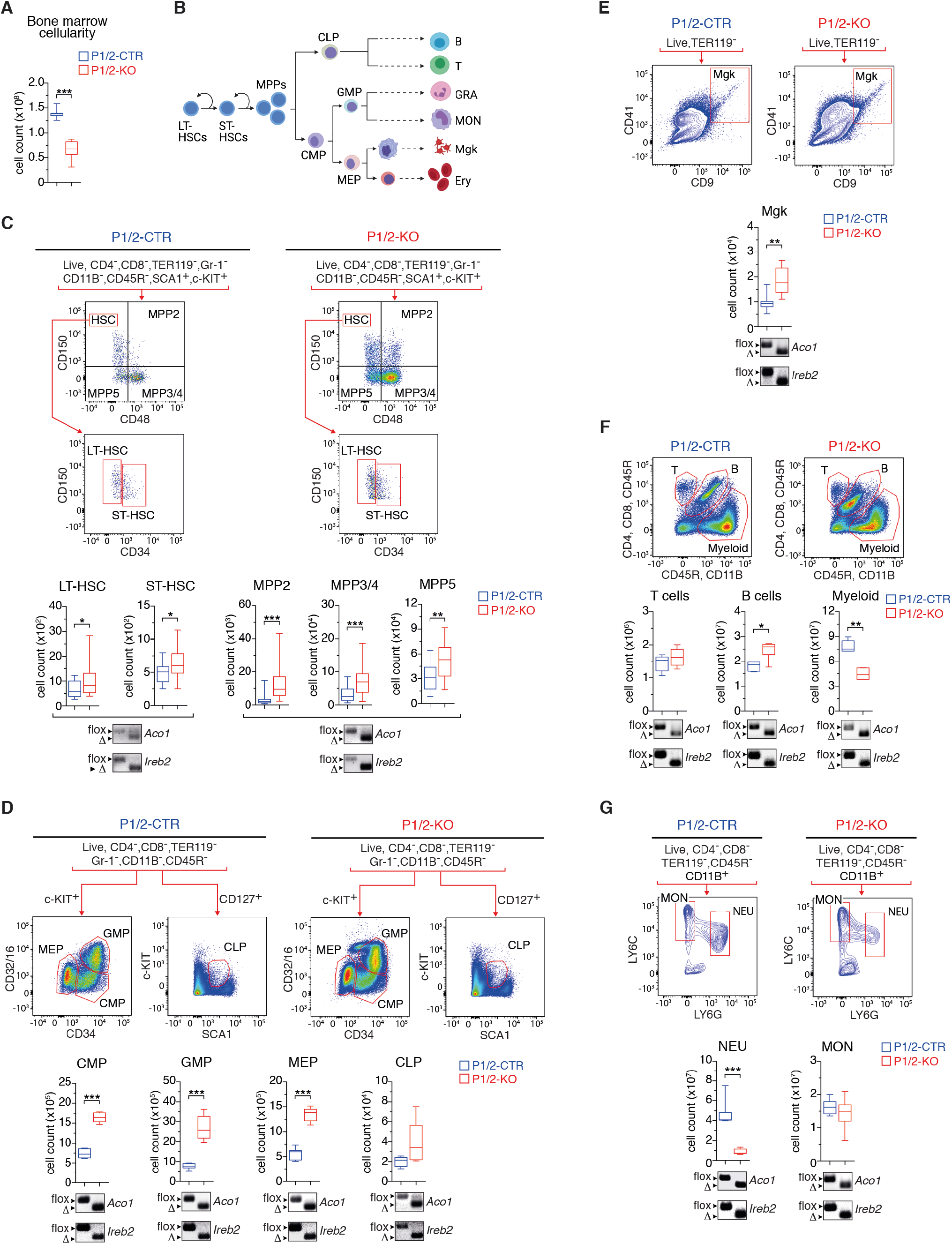
Immuno-phenotyping of bone marrow cell populations in adult mice with acute loss of IRP function. (A) Bone marrow (BM) cellularity. (B) Schematic representation of the hematopoietic system (Created with BioRender.com). (C-G) FCM analysis of BM cell populations: (C) long term (LT)- and short term (ST)-HSCs (hematopoietic stem cells) and multipotent progenitors (MPP, n=15-25); (D) common myeloid (CMP), lymphoid (CLP), megakaryocyte/erythroid (MEP), and granulocyte/macrophage (GMP) progenitors (n=5-10); (E) megakaryocytes (Mgk, n=5-9); (F) T- and B-cells, and myeloid cells (granulocytes/monocytes, n=8); (G) monocytes (MON) versus neutrophils (NEU) (n=7). The gating strategy is indicated for each cell population analyzed. Box plots (minimum to maximum values) show the number of cells in the BM of both hindlimbs. Genomic PCR panels below the box plots show recombination of the *Irp1* (*Aco1*) and *Irp2* (*Ireb2*) alleles in P1/2-KO (right) versus P1/2-CTR (left) mice (floxed: flox; truncated: Δ). *p<0.05; ** p<0.01; *** p<0.001. *See also Figure S2.*

Despite the expansion of lineage-negative (LIN^−^) cells, the number of differentiated erythroid (Figure 3A) and myeloid (Figure 2F) cells were substantially reduced. Consistent with the peripheral blood phenotype, there was an increase in megakaryocytes (Mgk, Figure 2E), and a slight elevation in B-cells (Figure 2F) in the BM. Hence, IRP ablation stimulates hematopoietic progenitor cells in the BM, while simultaneously suppressing erythroid and myeloid cells. Importantly, CRE activation alone did not alter BM cell populations (Figure S2G-L). To verify that the expanding LIN^−^ cells did not escape CRE recombination, we isolated stem and progenitor cells by FACS (fluorescence-activated cell sorting) and analyzed the recombination of the *Aco1* and *Ireb2* alleles by genomic PCR. Both alleles were efficiently recombined in all cell types tested (Figure 2C,D). *Irp* genes were also efficiently inactivated in Mgk as well as in B- and T-cells (Figure 2E,F).

**Figure 3:**
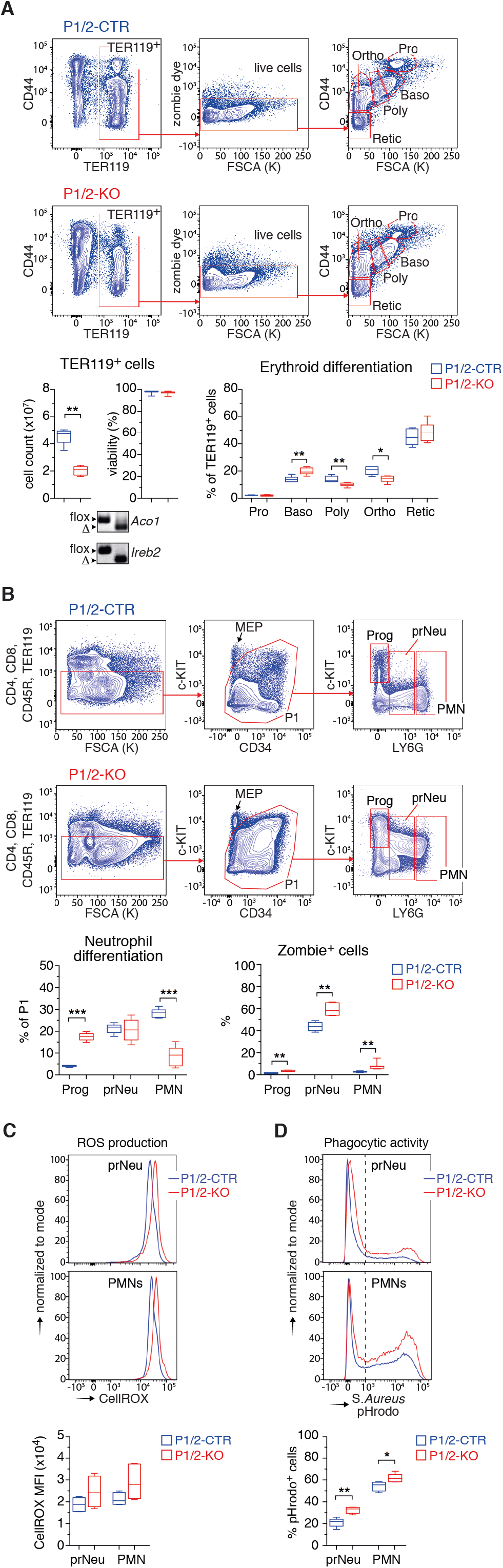
IRP deficiency impairs neutropoiesis. (A) FCM analysis of terminal erythroid differentiation (FSCA: forward side scatter area) in the BM (Pro: pro-erythroblasts; Baso, Poly, Ortho: baso, poly-, and ortho-chromatic cells, respectively; Retic: reticulocytes). Box plots (minimum to maximum values, n=5-6) show the number of TER119^+^ in the BM of both hindlimbs (left), the viability of TER119+ cells (middle), and the frequency of cells at different stages of erythroid differentiation (right). The panels below the box plots show the recombination of *Irp1* (*Aco1*) and *Irp2* (*Ireb2*) alleles in P1/2-KO (right) versus P1/2-CTR (left) mice (floxed: flox; runcated: Δ). (B) FCM analysis of neutrophil differentiation in the BM. c-KIT^high^LY6G^−^ represent progenitor (Prog) cells; c-KIT^−^LY6G^high^: polymorphonuclear (PMN) neutrophils; c-KIT^−^LY6G^low^: pre-neutrophils (prNeu). Box plots (minimum and maximum values, n=6-7) show the overall frequency of Prog, prNeu, and PMN cells in the BM (left), and the corresponding fraction of Zombie dye-positive cells (right). (C,D) FCM analysis of (C) ROS (reactive oxygen species) production and (D) engulfment of pHrodo-labeled S.*Aureus* particles. Top panels: representative FCM plots; bottom panel: data obtained from 5 mice (box plots with minimum to maximum values). In (D), the dotted line delimits pHrodo-positive and −negative cells. A-G: p-values correspond to separate pairwise comparisons between P1/2-CTR and P1/2-KO for each parameter and cell population analyzed (* p<0.05; ** p<0.01; *** p<0.001). *See also Figure S3.*

We next investigated whether the diminution of myeloid cells in the BM of P1/2-KO mice was confined to a specific subtype of cells. Despite efficient recombination of the floxed *Irp* alleles in both granulocytes and monocytes, IRP deficiency selectively suppressed neutrophils (defined as c-KIT^−^CD11B^+^LY6C^low^LY6G^high^ cells), which represent the major granulocyte subfraction (Figure 2G). Similarly to neutrophils, eosinophils were also significantly diminished to very low numbers, and basophils remained below the detection level (not shown). Monocytes (c-KIT^−^CD11B^+^LY6C^high^LY6G^−^) were unchanged (Figure 2G).

Overall, the IRP/IRE system does not appear to be strictly essential for the expansion of LIN^−^ hematopoietic progenitors in the BM, however it is required for the efficient production of erythroid cells and granulocytes.

### The IRP/IRE system is critical for neutrophil differentiation and survival in the bone marrow

We investigated the possible causes of the low number of erythrocytes and granulocytes in IRP-mutant mice. The survival of erythroid (TER119^+^) cells did not appear to be compromised (Figure 3A). Instead, we observed a defect in terminal erythroid differentiation, with impaired progression of erythroblasts from the basophilic to polychromatic stage. As erythroid differentiation is iron dependent (Muckenthaler et al., 2017), this defect could reflect a state of functional iron deficiency in P1/2-KO erythroblasts. Of note, CreER control animals exhibited normal erythropoiesis (Figure S3A).

Similarly, we examined the differentiation of neutrophils. c-KIT^high^LY6G^−^ progenitors (designated Prog) progressively lose the c-KIT marker and acquire LY6G expression as they differentiate into c-KIT^−^LY6G^high^ polymorphonuclear neutrophils (PMNs), with an intermediate stage (designated prNeu) characterized by intermediate LY6G expression and low to undetectable c-KIT levels (Figure 3B, top) (Riffelmacher et al., 2017). Consistent with the decrease in BM neutrophil counts (Figure 2G), P1/2-KO animals exhibited a marked decrease in PMN numbers, associated with an accumulation of Prog cells and the presence of immature prNeu cells positive for both c-KIT and LY6G (c-KIT^low^LY6G^high^) (Figure 3B). Importantly, CRE activation alone had negligible effects on neutrophil differentiation compared to IRP ablation (Figure S3B). The neutropoiesis defect in P1/2-KO mice was accompanied by a slight increase in the proportion of Prog cells in the S-phase of the cell cycle (Figure S3C). It was also associated with an augmentation of cell death (Figure 3B), which may contribute to the low abundance of LY6G^+^ cells.

We examined two of the key immune functions of LY6G^+^ neutrophils: their capacity to produce reactive oxygen species (ROS) and to engulf foreign particles upon stimulation with lipopolysaccharide (LPS, Borregaard, 2010, Németh, 2018). Surprisingly, IRP deficiency did not impair ROS production (Figure 3C), and in fact slightly increased the phagocytic activity of both prNeu and LY6G^high^ cells (Figure 3D).

Cumulatively, these results show that IRP deficiency impairs neutrophil differentiation and survival in the BM, yielding relatively immature neutrophils whose functionality appears to be preserved.

### IRPs promote neutropoiesis in a cell intrinsic manner

Physiological neutrophil differentiation could depend on IRP expression in cells of the hematopoietic niche or in the hematopoietic cells themselves. To distinguish between these two possibilities, we transplanted whole BM cells from WT mice into sub-lethally irradiated, untreated P1/2-KO recipients. Following stable engraftment, the resulting chimeras (designated WT→KO) were treated with tamoxifen to ablate the IRPs in non-hematopoietic tissues. As a control, WT BM cells were transplanted into P1/2-CTR recipients (WT→CTR). We found that IRP inactivation in non-hematopoietic cells does not alter neutrophil differentiation (Figure 4A). We next generated reverse chimeras where WT recipients were transplanted with BM cells from either P1/2-KO (KO→WT) or P1/2-CTR (CTR→WT) mice. Remarkably, KO→WT chimeras lacking IRP expression in hematopoietic cells reproduced the BM phenotype of P1/2-KO animals with systemic loss of IRP function. This included: 1) the reduction in RBC indices and concomitant increase in body iron, EPO, and hepcidin levels (Figure S4A-C), 2) the expansion of stem/progenitor cells (Figure S4D) and 3) the reduction in the number of PMNs in the BM, associated with the accumulation of c-KIT^high^ Prog cells and the presence of immature prNeus positive for both c-KIT and LY6G (Figure 4B). Taken together, the hematopoietic abnormalities and particularly the defects in neutropoiesis observed in P1/2-KO animals are BM intrinsic.

**Figure 4:**
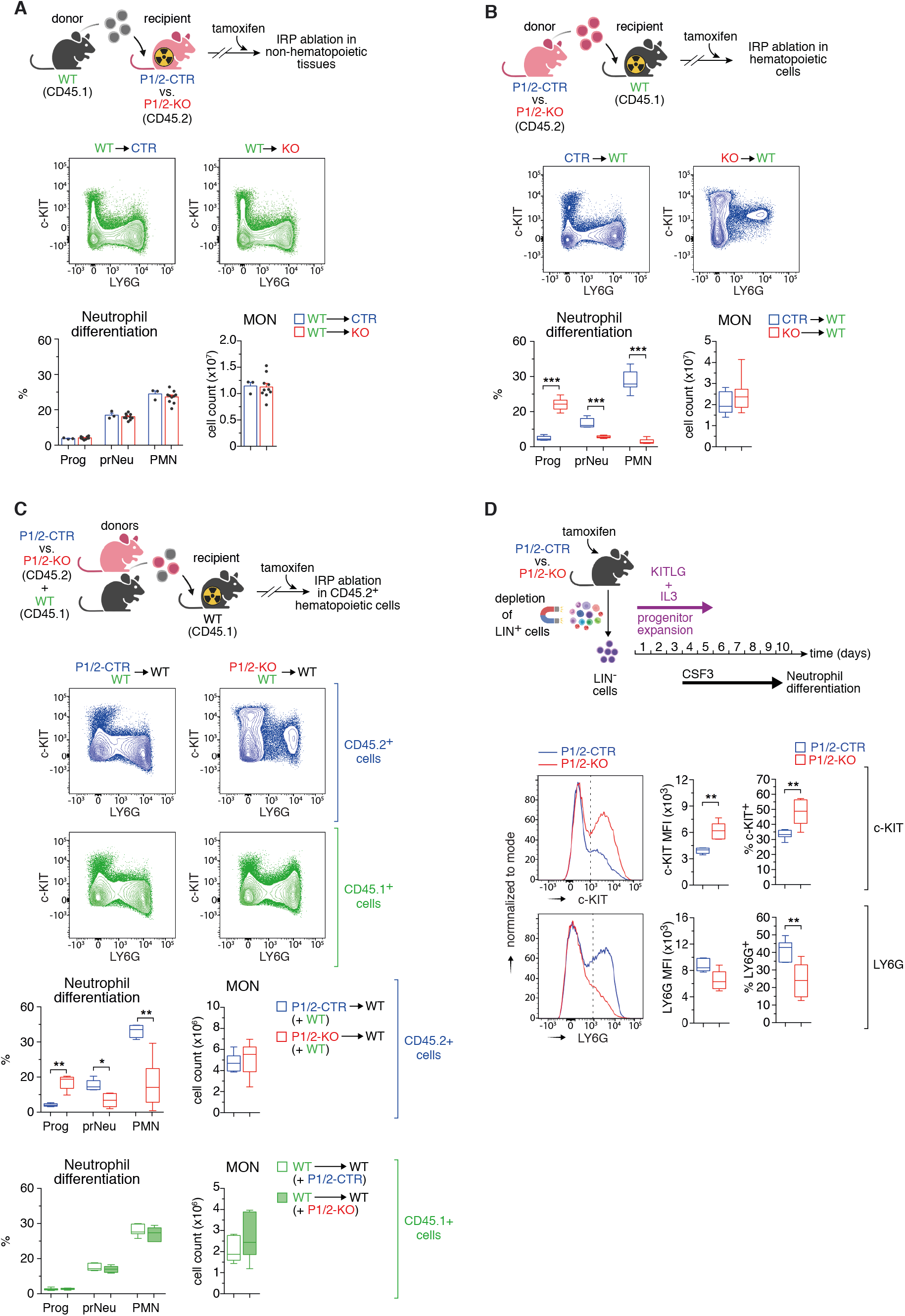
IRPs promote neutropoiesis in a cell-autonomous manner. (A, top) CD45.1^+^ BM cells from wild-type (WT) mice transplanted into irradiated P1/2-CTR versus P1/2-KO recipients. (B, top) CD45.2^+^ BM cells from P1/2-CTR versus P1/2-KO mice transplanted into WT recipients. (C, top) WT recipients transplanted with a 1:1 mixture of BM cells from P1/2-KO (versus P1/2-CTR) and WT mice. (A-C) Following stable engraftment, chimeras were treated with tamoxifen on day 1 and 3 and were analyzed on day 10. Representative FCM plots based on c-KIT and LY6G markers are show. Bar graphs and box plots display monocyte (MON) counts in the BM of both hindlimbs and the frequency of Prog, prNeu, and PMN cells (A, n=3-10; B, n=9-10; C, n=6-7). (D) Lineage-negative BM cells from tamoxifen-treated P1/2-CTR versus P1/2-KO mice were expanded *ex vivo* with KIT ligand (KITLG) and interleukin 3 (IL3), and then differentiated into LY6G^+^ neutrophils with CSF3. Bottom: Representative FCM plot analysis of c-KIT (top) and LY6G (bottom); the dotted line delimits marker-positive and −negative cells). Box plots (minimum to maximum values, n=7) display the median fluorescence intensity (MFI) for c-KIT and LY6G, respectively, and the percentage of cells positive for those markers. ** p<0.01; *** p<0.001. (A-D) top schemes created with BioRender.com. *See also Figures S4 and S5.*

To further investigate whether the neutropoietic phenotype of P1/2-KO mice is cell autonomous, we transplanted a 1:1 mixture of BM cells from WT mice (CD45.1^+^) and untreated P1/2-KO (CD45.2^+^) donors (or P1/2-CTR as control) into irradiated WT recipients (CD45.1^+^) and treated the mixed chimeras with tamoxifen. Hematopoietic cells derived from P1/2-KO donors show similar defects in neutropoiesis as those observed in P1/2-KO mice or KO→WT chimeras (Figure 4C). Furthermore, the presence of IRP-deficient cells did not affect the differentiation of precursor cells originating from WT donors (Figure 4C).

In addition, we exposed primary cultures of hematopoietic precursor cells from tamoxifen-treated P1/2-KO versus P1/2-CTR mice to G-CSF (granulocyte-colony stimulating factor, also CSF2) to mimic neutrophil differentiation *ex vivo* (Pelletier et al., 2017). Similar to the neutropoiesis defect seen *in vivo* in P1/2-KO mice, we observed an increase in c-KIT levels and an accumulation of c-KIT^+^ cells to the detriment of LY6G^+^ cells (Figure 4D). Of note, the effect of IRP deficiency on cell development seems neutrophil lineage specific, as monocyte counts in the BM of transplanted mice are unaffected (Figure 4 A-C) and the differentiation of P1/2-KO precursors into LY6C^+^ monocytes *ex vivo* is normal (Figure S5A).

Together, these results demonstrate that IRPs are required cell-intrinsically for neutropoiesis.

### IRPs are required to complete terminal neutrophil differentiation in the bone marrow

To better understand how IRP deficiency alters neutrophil differentiation, we analyzed the transcriptome of Prog, prNeu and PMN cell populations isolated from P1/2-KO versus P1/2-CTR mice. Most changes in gene expression can be accounted for by the differentiation stage of the cell, regardless of the IRP status (Figure 5A). Nonetheless, IRP-deficient cells cluster separately from control at all stages of differentiation (Figure 5A). We compared Prog, prNeu, and PMN cell populations in either P1/2-CTR or P1/2-KO mice and identified the differentially expressed genes (DEGs, Table S1). DEGs were then grouped in 4 main clusters (Table S2) reflecting their overall expression dynamics in P1/2-CTR cells (Figure 5B). Consistent with the acquisition of specialized cellular functions (Grassi et al., 2018), a majority of DEGs were either down- (cluster 1) or up- (cluster 3) regulated during neutropoiesis (Figure 5B,C). Other genes exhibited a transient decrease (cluster 2) or peak (cluster 4) in expression at the intermediate differentiation stage (Figure 5B). Importantly, the expression pattern of genes coding for transcription factors required for myeloid development (Evrard et al., 2018) were very similar in P1/2-KO and P1/2-CTR cells (Figure S6A). This included CEBPA (CCAAT Enhancer Binding Protein α), which is needed for the generation of GMPs from HSCs, as well as CEPBE and GFI1 (Growth Factor Independent 1 Transcriptional Repressor), both of which are important for neutrophil development (Lawrence et al., 2018, Ai and Udalova, 2020). Likewise, the expression trajectory of genes typically associated with neutrophil differentiation, such as those encoding the antimicrobial factors contained in azurophilic (AGs), specific (SGs), and gelatinase (GGs) granules, was largely preserved in P1/2-KO cells (Figure S6A). This suggests that the transcriptional programs driving neutropoiesis are not completely disrupted in IRP deficiency.

**Figure 5:**
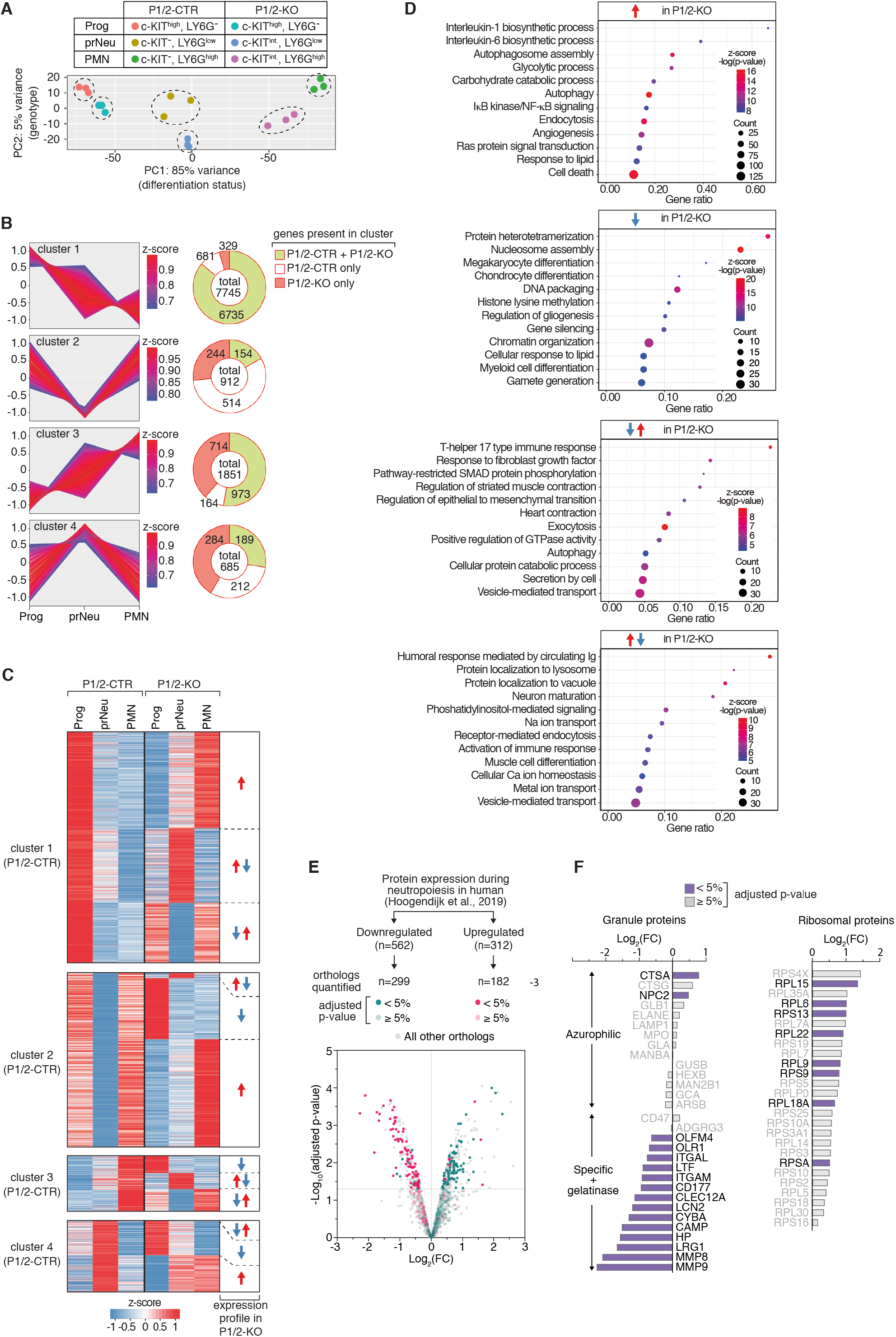
Impact of IRP deficiency on gene expression dynamics during neutropoiesis. (A-D) RNA-seq analysis of the transcriptome of Prog, prNeu and PMN cell populations isolated from the BM of P1/2-CTR versus P1/2-KO mice. (A) Principle component (PC) analysis. (B) Genes differentially expressed during neutropoiesis were grouped in 4 main categories using k-means clustering (left). The pie charts (right) indicate: 1) the total number of genes in a cluster, 2) the number of genes found in that cluster only in P1/2-CTR (white) or in P1/2-KO (orange) cells, respectively, and 3) the number of genes with similar expression patterns in both P1/2-CTR and P1/2-KO cells (green). (C) Heatmaps display expression profiles that differ between P1/2-CTR and P1/2-KO cells. For each cluster in P1/2-CTR cells, the expression trajectory of the corresponding genes in P1/2-KO cells are shown. (D) Gene ontology enrichment analysis of genes dysregulated during neutropoiesis in P1/2-KO cells. The dot plots show enrichment for biological processes. (E) Proteome analysis of whole BM LY6G^+^ cells from P1/2-KO versus P1/2-CTR mice. The colors highlight proteins whose orthologs have been reported to be either down- or up-regulated during neutrophil differentiation in human BM (Hoogendijk et al., 2019). FC: fold change, P1/2-KO – P1/2-CTR. (F) Bar graphs: FC expression for selected granule (left) and ribosomal (right) proteins. In (E) and (F), the color code indicates the p-value adjusted with the Benjamini–Hochberg method for multiple testing. *See also Tables S2-S4, and Figure S6.*

16% of all DEGs displayed a distinct expression pattern in P1/2-KO cells when compared to P1/2-CTR (Figure 5B and 5C). Among those 1571 genes, nearly half (714) were upregulated in P1/2-KO cells only, whereas 329 were downregulated (Figure 5B). qRT-PCR analysis of selected transcripts confirmed these expression profiles and also showed that those molecular features were due to IRP deficiency and not mere consequences of CRE activation (Figure S6B). Of note, the gene expression profiles in P1/2-CTR cells did not predict expression profiles in IRP-mutant cells. That is, genes belonging to a given cluster of P1/2-CTR cells were found to cluster in any of the other three groups of mutant cells (for example, the expression of genes normally upregulated during neutrophil development may either increase or transiently peak/drop in IRP deficiency) (Figure 5C and Table S2). A GO (gene ontology) enrichment analysis (Table S3) revealed that genes displaying a transient increase (n=284) or decrease (n=244) in expression in P1/2-KO cells were related to cellular vesicles biology, secretion, immune responses, signal transduction, and cytoskeleton organization (Figure 5D and Table S3). mRNAs whose expression is abnormally downregulated in IRP deficiency (n=329) were mainly involved in cell differentiation programs and chromatin organization (Figure 5D and Table S3). Changes in chromatin architecture play a key role in the regulation of neutrophil differentiation genes (Zhu et al., 2017, Manley et al., 2018), suggesting that these alterations in gene expression may reflect the inability of P1/2-KO cells to undergo normal neutrophil differentiation. Transcripts displaying increased expression in IRP deficiency (n=714) encoded, among others, proteins involved in glycolysis and autophagy, as discussed later.

To complement our transcriptome study and assess how mRNA dysregulation in IRP-deficient neutrophils affects protein expression, we analyzed the proteome of total LY6G^+^ BM cells from P1/2-KO and P1/2-CTR mice. We focused on the 1942 proteins that yielded a signal in 70% of the samples in at least one of the genotypes analyzed, and could thus be quantified with high confidence (Table S4). We cross-referenced our data with a published dataset describing the proteomic changes that accompany the differentiation of pre-myelocytes towards mature neutrophils in humans (Hoogendijk et al., 2019). Of the 312 proteins reported to be upregulated during neutropoiesis in humans, 182 mouse orthologs are present in our dataset. Interestingly, most of these 182 proteins were less abundant in Ly6G^+^ cells from P1/2-KO mice (Figure 5E). These proteins were mainly constituents of the SGs/GGs (Figure 5F) that mark the final stages of neutrophil development in the BM (Rørvig et al., 2013, Hoogendijk et al., 2019). In comparison, the abundance of AG protein markers expressed during early neutropoiesis remained unchanged, suggesting that IRP deficiency impairs intermediate stages of neutrophil development. Conversely, the majority of the 312 proteins whose human orthologs were suppressed during neutropoiesis showed a trend towards higher expression in IRP deficiency (Figure 5E and Table S4). Several of these proteins are components of the ribosome (Figure 5F). Biosynthetic processes usually decline as cells acquire neutrophil functions (Hoogendijk et al., 2019). As such, the elevation of ribosomal proteins in LY6G^+^ cells from P1/2-KO mice is an additional sign that neutrophil differentiation is incomplete.

Overall, IRP deficiency does not seem to disrupt the core transcriptional program that drives neutropoiesis. Our data rather indicate that IRPs act in parallel with the gene regulators governing neutrophil development to facilitate the production of fully differentiated neutrophils in the BM.

### IRP deficiency impairs metabolic rewiring during neutrophil development

A substantial fraction of mRNAs dysregulated in IRP deficient cells are abnormally stimulated (n=714) during neutrophil development (Figure 5B,C and Table S2). These transcripts encode proteins mainly associated with GO-terms related to autophagy, glucose catabolism, and cell death (Figure 5D and Table S3). The upregulation of cell death-related genes is consistent with the increased frequency of Zombie dye^+^ cells in P1/2-KO mice (Figure 3B). However, the upregulation of genes involved in glycolysis and autophagy was unexpected, as both processes have been shown to decline during neutrophil development in the BM (Kumar and Dikshit, 2019, Riffelmacher et al., 2017). We further interrogated our proteomics data based on the idea that the protein level of rate-limiting enzymes determines the activity of metabolic pathways in the cell (Ahl et al., 2020). Indeed, LY6G^+^ cells from P1/2-KO mice expressed high levels of the phosphofructokinase isoenzymes (PFKL, PFKP), which promote glycolysis (Tanner et al., 2018), and downregulated anti-glycolytic enzyme fructose bisphosphatase (FBP1, Figure 6A). Furthermore, we observed that the uptake of the 2-NBDG glucose analogue during neutrophil development was higher in P1/2-KO mice than in P1/2-CTR mice; further evidence for increased glycolytic flux (Figure 6B). Similarly, FCM analysis of autophagic vacuole formation confirmed the augmentation of autophagic activity in LY6G^+^ cells of IRP-mutant animals (Figure S6C). We also noticed “response to lipids” as a GO term associated with genes that are either up or down-regulated in IRP deficiency (Figure 5D and Table S3), and observed vacuoles resembling lipid droplets in P1/2-KO myelocytes (not shown). FCM analysis with the BODIPY 493/503 dye revealed the accumulation of neutral lipids at all stages of neutrophil differentiation (Figure S6D). This suggests that IRP deficiency may impair lipid metabolism or transport, or trigger lipid droplet formation as a protective measure (Olzmann and Carvalho, 2019).

**Figure 6:**
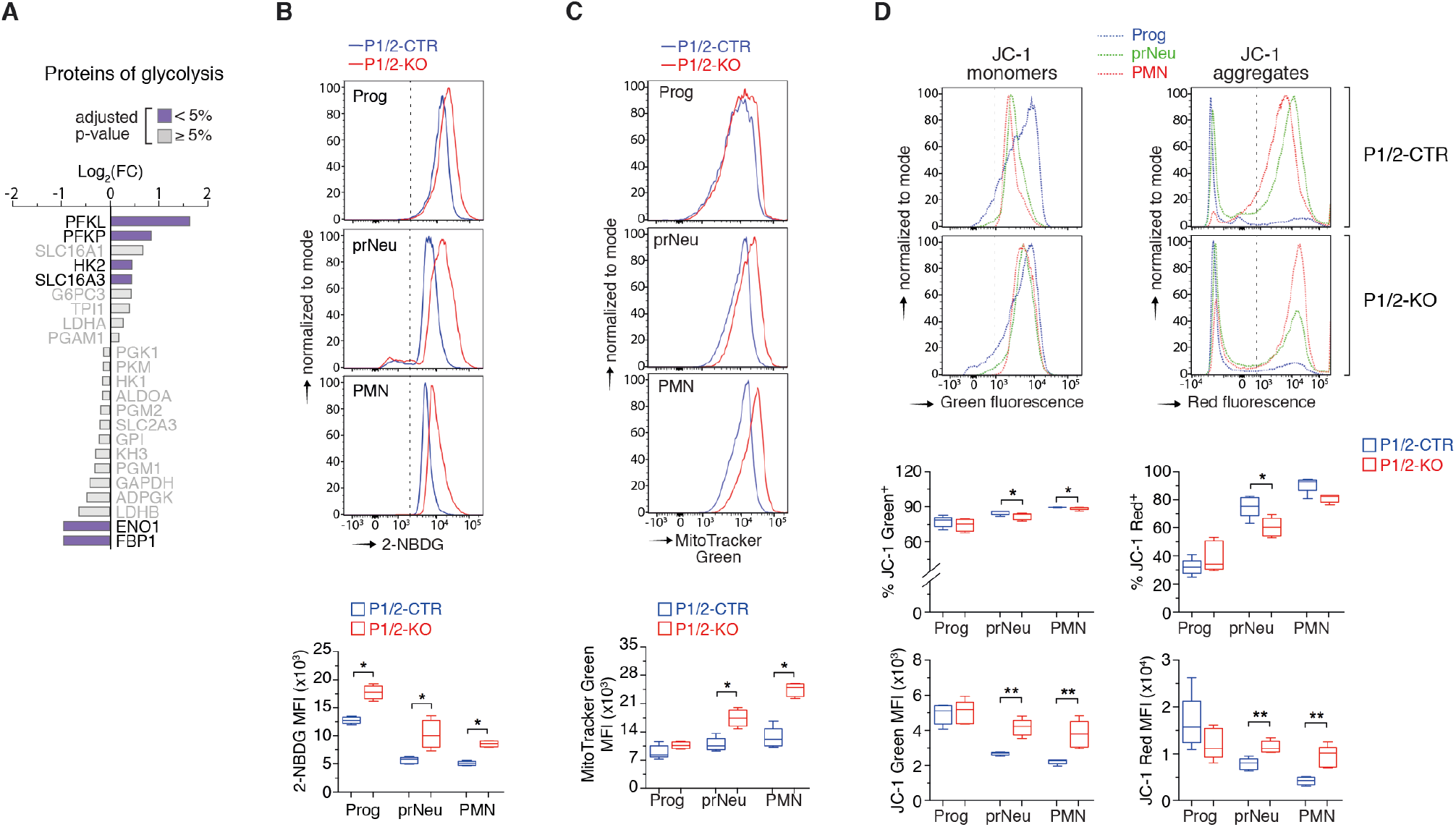
IRP deficiency causes abnormal metabolic remodeling during neutrophil development and differentiation. (A) Expression of selected glycolysis proteins in whole BM LY6G^+^ cells (same representation as Figure 5F). (B-D) FCM analysis of (B) 2-NBDG uptake, (C) mitochondrial mass (with MitoTracker green dye), and (D) mitochondrial membrane potential (with JC-1 dye) during neutropoiesis. Top: representative FCM plots (the dotted line separates stained from unstained cells). Bottom: box plot with maximum to minimum MFI values (A, n=4-5; B,E n=5). Middle box plot in (D): % of JC-1 positive cells (maximum to minimum values, n=5). p-values correspond to separate pairwise comparisons between P1/2-CTR and P1/2-KO for each parameter and cell population analyzed. * p<0.05; ** p<0.01. *See also Figure S6 and Tables S4 and S5.*

Mitochondria in fully differentiated blood neutrophils are present in small numbers and serve to control cell survival rather than to produce energy (Hoogendijk et al., 2019). However, in the BM, adequate neutrophil differentiation is dependent on a metabolic shift that limits glycolysis while increasing mitochondrial content and respiration (Riffelmacher et al., 2017). Mitochondria use iron to produce energy via heme- and ICS- proteins (Ward and Cloonan, 2019) and mitochondrial integrity has been shown to be compromised in IRP-deficient hepatocytes (Galy et al., 2010). We therefore hypothesized that IRP ablation might alter neutropoiesis by reducing the content and/or activity of mitochondria in hematopoietic cells, resulting in a decrease in mitochondrial proteins. Our proteomics data revealed a strong enrichment of GO terms related to the cellular component mitochondria, suggesting changes in mitochondrial protein abundance (Table S5). Of the proteins listed in the mouse MitoCarta database (Rath et al., 2021), 222 could be quantified in LY6G^+^ cells. Interestingly, most of these proteins were expressed at a higher level in P1/2-KO cells compared to P1/2-CTR cells and accounted for as much as one-fourth of all proteins significantly upregulated in IRP deficiency (Figure S6E and Table S4). In contrast, only 2 mitochondrial proteins were significantly reduced in P1/2-KO cells (4% of all downregulated proteins). Hence, contrary to expectations, the mitochondrial content of LY6G^+^ cells appeared to be increased in IRP deficiency. Accordingly, FCM analysis with MitoTracker green dye revealed that the mitochondrial mass is markedly augmented in prNeu and PMN cells from P1/2-KO mice (Figure 6D).

To test whether IRP deficiency could lead to the accumulation of unhealthy mitochondria, we analyzed the mitochondrial membrane potential (MMP) using the mitochondrial dye JC-1. JC-1 monomers emit green fluorescence in cells with low MMP, but JC-1 forms aggregates producing red fluorescence in cells with high MMP; changes in the intensity of red versus green signal reflect changes in the cell’s MMP (Cossarizza et al., 2019). In P1/2-CTR mice, the proportion of JC-1 red^+^ cells increased from the Prog to PMN stage, while the ratio of red to green signal intensity is globally preserved (Figure 6C). Similarly, the percentage of JC-1red^+^ cells in P1/2-KO mice increased during neutropoiesis, albeit to a slightly lesser extent at the prNeu stage (Figure 6C). Both red and green signal intensities were elevated in prNeu and PMN cells (Figure 6C), possibly reflecting the expansion of mitochondrial mass and corresponding increase in JC-1 dye uptake. However, we did not detect a significant gain in green versus red fluorescence intensity, which could have indicated membrane depolarization. Acute IRP ablation thus appeared to increase the amount of apparently healthy mitochondria during neutropoiesis.

Together, these data suggest that IRP deficiency interferes with the metabolic remodeling that accompanies neutrophil development and differentiation in the BM, possibly reflecting a situation of stress characterized by high glycolytic and autophagic activities, lipid accumulation, and expansion of the mitochondrial content of the cell.

### The IRP/IRE network supports neutropoiesis by securing iron bioavailability

A transcriptome-wide search for IRP-interacting mRNAs (Sanchez et al., 2011) and recent *in cellulo* RNA foot-printing data (Corley et al., 2020) suggest that the IRP regulome extends beyond iron metabolism genes. This raised the question of whether the neutropoiesis defect of IRP-deficient mice is caused by alterations in iron metabolism or iron-independent IRP functions. We therefore examined how IRP deficiency affects the cellular iron status during neutropoiesis. IRPs normally inhibit the decay of TFRC mRNA and hamper the translation of FTL1, FTH1, and FPN transcripts. Accordingly, Prog cells from P1/2-KO mice exhibited a strong decrease in TFRC expression compared with P1/2-CTR; TFRC levels in LY6G^high^ cells remained below detection regardless of genotype (Figure 7A). Conversely, ferritin expression was higher in IRP deficiency in both LIN^−^ and LY6G^+^ cells (Figure 7B). FPN expression was elevated in LY6G^+^ cells, but not in progenitor cells (Figure 7C). Downregulation of TFRC together with stimulation of the ferritin iron sequestering molecules and FPN iron exporter (in LY6G^+^ cells) is predicted to lower cellular iron availability.

**Figure 7:**
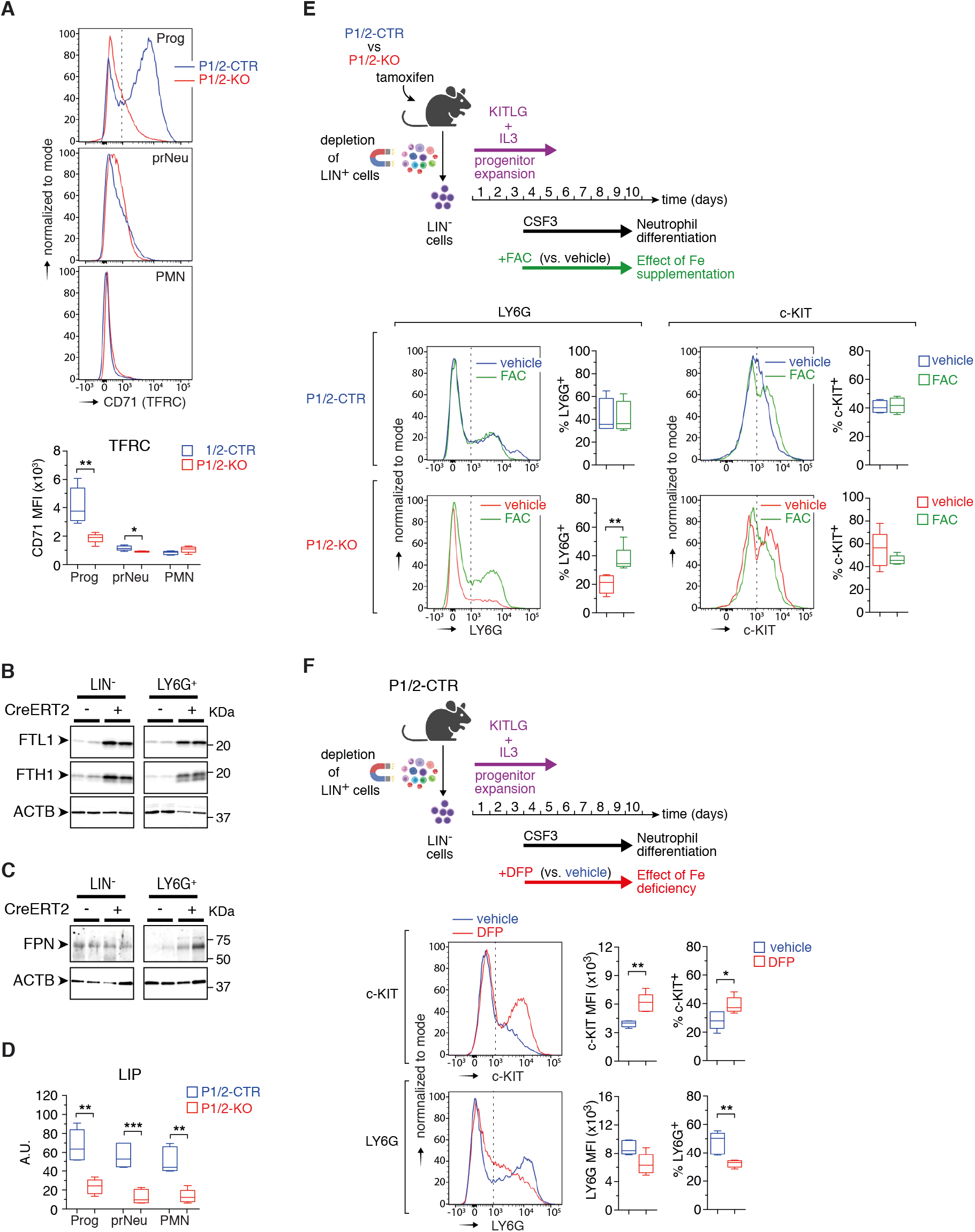
IRPs support neutrophil differentiation by securing iron bioavailability. (A) FCM analysis of CD71/TFRC expression during neutropoiesis *in vivo*. Top: representative FCM plots. Bottom: box plot with maximum to minimum MFI values (n=5-6). (B,C) Representative western-blot analysis of ferritin (B) and FPN (C) expression in lineage negative (LIN^−^) versus LY6G^+^ cells from the BM of P1/2-CTR (CreERT2-) versus P1/2-KO (CreERT2+) mice. Loading control: ACTB (actin ß) (D) Box plot (maximum to minimum values) showing reduced labile iron pool (LIP) in BM cells of P1/2-KO versus P1/2-CTR mice during neutropoiesis *in vivo* (n=5-6). (E,F) Top schemes (created with BioRender.com): *Ex vivo* differentiation of LIN^−^ BM cells as in Figure 4D, in the presence of either (E) ferric ammonium citrate (FAC) or (F) deferiprone (DFP); control cells were treated with vehicle. Bottom: representative FCM plots showing LY6G and c-KIT levels, together with box plots (maximum to minimum values, n=4-5) displaying the MFI (F) or % of cells positive (E,F) for either marker. The dotted lines delineate marker-positive and −negative cells. (A,D-F) p-values correspond to separate pairwise comparisons between P1/2-CTR and P1/2-KO for each cell population and/or parameter analyzed. * p<0.01; ** p<0.01; *** p<0.001. *See also Figure S7.*

P1/2-KO BM cells showed a marked reduction in the metabolically active labile iron pool (LIP) during neutropoiesis compared to P1/2-CTR cells (Figure 7D). A similar reduction in the LIP was observed in P1/2-KO cells undergoing *ex vivo* differentiation (not shown). These observations prompted us to test whether an extra source of iron could restore normal neutropoiesis. For this, we used our *ex vivo* neutrophil differentiation assay and exposed cells from tamoxifen-treated P1/2-KO mice to ferric ammonium citrate (FAC), a form of iron that cells can take up independently of TFRC (Figure 7E, top). Compared to vehicle, P1/2-KO cells treated with 10 µg/mL of FAC differentiated more efficiently, whereas the same amount of FAC had no effect on cells from P1/2-CTR mice (Figure 7E). Another iron donor, iron sulfate pentahydrate, had a similar effect as FAC (not shown). These results strongly suggest that IRP ablation impairs neutropoiesis due to iron shortage.

As a complementary approach to the iron rescue experiment, we exposed cells from untreated P1/2-CTR mice to the iron chelator deferiprone (DFP; Figure 7F, top). DFP treatment mimicked the effect of IRP ablation (Figure 4D), with stimulation of c-KIT expression and accumulation of c-KIT^+^ cells at the expense of LY6G^+^ cells (Figure 7F). In contrast to neutrophils, DFP stimulated the differentiation of hematopoietic progenitors towards LYC6^+^ monocytes (Figure S5B). This indicates that neutrophil differentiation is particularly sensitive to iron deprivation. *In vivo*, chronic treatment with Ferriprox, a DFP oral formulation, had only a minor effect on steady-state neutropoiesis (Figure S7A-E). This could potentially be explained by relatively modest iron deprivation in BM cells compared to other cells, as the compound appears to have uneven effects on iron. Indeed, DFP could deplete the liver iron store very effectively, but had no measurable effect on spleen iron (Figure S7B). Nevertheless, Ferriprox altered neutrophil differentiation when mice were additionally treated with CSF2/G-CSF (in the form of Filgrastim/Neupogen) to mimic conditions of emergency granulopoiesis, with a significant increase in c-KIT levels (Figure S7C,D) and accumulation of immature prNeu cells at the expense of LY6G^high^ PMN cells (Figure S7E). This reinforces the notion that iron bioavailability influences neutrophil differentiation in the BM.

## DISCUSSION

The majority of iron in the human body is contained in hemoglobin and dedicated to oxygen transport by erythrocytes (Pasricha et al., 2021). While the importance of iron for RBC formation is undisputed, its role in other facets of blood cell biology is only beginning to be recognized, with notably recent reports showing a role for the metal in adaptive immunity (Jabara et al., 2016, Jiang et al., 2019, Frost et al., 2021). Through genetic disruption of a key iron homeostatic machinery in mice, our work revealed that maintenance of cellular iron bioavailability is critical for granulocyte development and differentiation in the BM, establishing a previously under-appreciated link between iron metabolism and a major arm of the innate immune system.

Studies in mice demonstrated that the IRP/IRE regulatory network is essential during early embryonic development (Smith et al., 2006). The hepatocytic and intestinal functions of the IRP/IRE system are also essential during perinatal life (Galy et al., 2008, 2010), but seem dispensable during adulthood in unchallenged animals (unpublished observations). Moreover, constitutive loss of both IRP1 and IRP2 in the heart or macrophages, respectively, remains asymptomatic under standard laboratory conditions (Haddad et al., 2017, Nairz, Ferring-Appel et al., 2015). These observations suggest that the IRP/IRE system exerts particularly important functions during pre- and peri-natal life. Here, we used a temporally-controlled CRE system enabling acute disruption of IRP1 and/or IRP2 in nearly the entire body of adult mice. Two evident short-term manifestations of acute IRP ablation in adult mice were microcytic anemia and neutropenia. The neutropenia was not observed in mice harboring a single loss of either IRP1 or IRP2, therefore reflecting overlapping functions between the two proteins. The anemia, on the other hand, was attributable to IRP2 deficiency and corroborates previous findings in standard mouse models with constitutive loss of IRP2 function (Copperman et al., 2005, Galy et al., 2005b). In this study, we focused on the role of the IRPs in hematopoiesis, but acute IRP deficiency may have additional effects yet to be investigated. Future work may uncover a role for the IRP/IRE network in other organs, at different stages of adult life (e.g. during ageing), or in response to challenge.

Because of the importance of iron in oxygen transport, research on iron metabolism has focused primarily on the biology of RBCs. The role of the metal in overall hematopoiesis and, in particular, HSCs remains poorly understood. Iron homeostasis can affect HSCs in different manners. Iron overload due to ineffective erythropoiesis or RBC transfusion in patients with hematological diseases is known to impair HSC functions (Weber et al., 2021). Iron excess can be detrimental to cells of the hematopoietic niche (Lu et al., 2013, Tanaka et al., 2019); it can also damage the HSCs themselves, as illustrated by the stem cell exhaustion phenotype of FBXL5-null mice with selective iron accumulation in HSCs (Muto et al., 2017). However, the impact of iron deficiency on HSCs is less clear. Iron deprivation with DFO diminished the proliferation and survival of hematopoietic progenitors in an *in vitro* model of mouse embryonic hematopoiesis (Shvartsman et al., 2019). Whereas iron chelation in human and mouse HSCs with Eltrombopag was shown to stimulate stem cell self-renewal and multilineage hematopoiesis (Kao et al., 2018). Our study revealed that hematopoietic stem/progenitor cells can expand in the absence of a fully functional IRP/IRE system and withstand the ensuing cellular iron deficiency, at least over a short period of time. Although we cannot exclude that IRP ablation may impair the stem/progenitor cell compartment in the long term, these data suggested that the iron requirements of hematopoietic stem/progenitor cells are relatively modest. Similarly, IRP-mediated iron homeostasis was dispensable for lymphoid development in the upper part of the hematopoietic hierarchy. This may reflect the fact that the flow of multipotent progenitors to the lymphoid lineage is low compared to the erythro-myeloid lineage, as shown by *in vivo* fate mapping (Höfer et al., 2016). This does not imply, however, that iron is not important for lymphoid compartment expansion at later stages of differentiation and, indeed, iron availability has been shown to be critical during lymphocyte activation (Frost et al., 2021, Teh et al., 2021).

Both RBCs and platelets are derived from MEPs. While IRP ablation suppressed RBCs, it caused an increase in platelet counts. The expansion of Mgk in the BM of IRP-mutant mice may reflect increased commitment of MEPs towards the Mgk lineage in conditions of iron deficiency (Xavier-Ferrucio, Scanlon et al., 2019). Mgk might also bypass the MEP progenitors and arise directly from the enlarged MPP2 population (Psaila and Mead, 2019).

The most profound and unexpected effect of acute IRP ablation was the significant decrease in neutrophis. This effect was in part due to functional iron deficiency in hematopoietic cells. While iron overload and high heme levels are known to impair the function of mature neutrophils (Martins et al., 2016, Renassia et al., 2020), how iron scarcity specifically affects neutrophils is not well understood (Abuga et al., 2020). Cases of neutropenia associated with iron deficiency and reversed upon iron therapy have been reported (Abuirmeileh et al., 2014, Abdelmahmuod et al., 2020). Also, DFP, an iron chelator mainly used to treat iron overload in Thalassemia patients, has been shown to cause neutropenia via a yet unclear mechanism (Kontoghiorghes et al., 2020). Although the cases are rare, such a side effect can increase the risk and severity of infection (Tricta et al., 2016). Accordingly, we found that DFP limits neutrophil differentiation *ex vivo*. *In vivo*, DFP could alter neutrophil development upon pharmacological stimulation of neutropoiesis with G-CSF. Under the experimental conditions used in our study, the effect of DFP in mice was rather mild compared to the profound granulopoiesis defect seen in IRP-mutant animals, possibly due to insufficient access of the compound to the iron pool of BM cells.

IRP ablation suppressed granulocytes *in vivo* but did not reduce the abundance of monocytes, the other major constituent of the myeloid compartment. This was remarkable because both cell types are derived from GMPs, are relatively short-lived, and reside only briefly in the BM (Yona et al., 2013). Interestingly, we found that DFP promotes the differentiation of hematopoietic stem/progenitor cells into monocytes *ex vivo*. This is consistent with previous observations showing a positive effect of iron deprivation on the differentiation of cord blood cells and leukemic cell lines towards the monocytic lineage (Callens et al., 2010). Iron deficiency thus impairs granulopoiesis but at the same time appears to favor monopoiesis, suggesting that iron bioavailability may be critical for shaping the innate defense system. The differential effect of iron on granulopoiesis versus monopoiesis and which specific differentiation node is modulated by iron availability will be an exciting question to explore.

Iron metabolism is not needed to initiate neutropoiesis *per se*, as the expression trajectory of granule genes and transcription factors important for neutrophil differentiation is globally preserved in IRP deficiency. Iron rather operates in parallel to the gene regulators that drive neutropoiesis to enable the production of fully differentiated neutrophils. A similar role of metabolism has been reported for autophagy. Differentiating neutrophils mobilize cellular lipids through lipophagy to fuel fatty acid oxidation, thereby promoting a shift from glycolysis to oxidative phosphorylation that is critical for neutrophil differentiation (Riffelmacher et al., 2017). IRP ablation led to high glycolytic and autophagic activity, lipid droplet accumulation, and increased mitochondrial content, indicating that the metabolic rewiring that accompanies neutrophil differentiation cannot operate normally in IRP deficiency. Among other possibilities, elevated glycolysis and high mitochondrial content could be a compensatory response to an energy stress, whereas lipid droplets could form because of defective fatty acid oxidation or to avoid lipotoxicity during autophagy (Olzmann and Carvalho, 2019). To what extent these metabolic alterations may contribute to the neutropoietic defect of P1/2-KO mice, and how they are linked to iron metabolism, is not clear. Importantly, hundreds of proteins in the cell bind iron either as heme, ISC or iron ions (Andreini et al., 2018), and hence multiple iron-dependent pathways may act together to support neutropoiesis. Future research should aim at deciphering the exact mechanism through which IRPs and iron promote neutrophil differentiation and survival in the BM.

Overall, our study reinforces the notion that iron metabolism influences multiple aspects of hematopoiesis and plays important roles in shaping the immune system. We showed that IRP-mediated iron homeostasis is critical for neutrophil development in the BM. Such finding could be relevant, for instance, in the context of infection/inflammation-induced hypoferremia. This process of iron sequestration in iron stores, which is considered a mean for the host to restrict access of iron to circulating siderophilic pathogens, can limit the supply of iron to erythroblasts and cause anemia (Haschka et al., 2021). Our study suggests that acute inflammatory hypoferremia may also alter neutrophil development in the BM, which could potentially compromise host defenses. The influence of iron on neutropoiesis may also have implications for G-CSF-based therapies in patients receiving myelosuppressive antineoplastic treatments or BM transplants, as our study suggests that the iron status of the patient and/or transplanted cells may impact the efficacy of the treatment.

## ACKNOWLEDGEMENTS

We thank the Center for Preclinical Research as well as the Flow Cytometry and the Genomics and Proteomics core facilities of the DKFZ for their support. We wish to thank Matthias W. Hentze (EMBL) for sharing the floxed *Irp* mouse lines, and A. Ruggieri (Heidelberg University) for her critical reading of the manuscript. This work was supported by a grant from the DFG (GA 2075/6-1) to B.G.

## AUTHOR CONTRIBUTION

Designed the research: MB and BG.

Performed experiments: MB, ET, CK, MQ, GP, MPS, WR, AE, NA, BG.

Analyzed the data: MB, SA, MS, DH, RB, MDM, BG.

Wrote and reviewed the manuscript: MB and BG with the help of all co-authors.

