## Supplemental for "IRON REGULATORY PROTEIN (IRP)-MEDIATED IRON HOMEOSTASIS IS CRITICAL FOR NEUTROPHIL DEVELOPMENT AND DIFFERENTIATION IN THE BONE MARROW"

- **STAR*methods including key resources table**

- Figure S1: **Acute ablation of either IRP1 or IRP2 does not impair steady-state myelopoiesis,** related to Figure 1.
- Figure S2: **Effect of CRE activation on blood cell counts, serum iron parameters, and bone marrow cell populations**, related to Figures 1 and 2.
- Figure S3: **Effect of CRE on erythroid and myeloid differentiation in the bone marrow, and consequences of IRP deficiency on the cell cycle during neutropoiesis**, related to Figure 3.
- Figure S4: **Mice lacking IRP expression in hematopoietic cells phenocopy mice with systemic loss of IRP function**, related to Figure 4.
- Figure S5. **Effects of IRP deficiency and iron deprivation on the differentiation of hematopoietic progenitor cells towards monocytes**, related to Figures 4 and 7.
- Figure S6: **Validation of the analysis and interpretation of transcriptome and proteome data**, related to Figure 5.
- Figure S7: **Effect of chronic treatment with the iron chelator deferiprone on steady-state and emergency neutropoiesis *in vivo***, related to Figure 7.

- Table S1: **RNA-seq data, genes differentially expressed during neutropoiesis in P1/2-CTR and P1/2 -KO mice (analysis of sorted Prog, prNeu and PMN cell populations)**, related to Figure 5.
- Table S2: **Clustering of genes differentially expressed during neutropoiesis in P1/2-CTR versus P1/2 -KO mice (based on RNA-seq data)**, related to Figures 5.
- Table S3: **GO enrichment analysis of mRNAs misregulated during neutropoiesis in P1/2-KO mice (based on RNA-seq data)**, related to Figures 5, 6, and S6.
- Table S4: **Proteomics data, analysis of LY6G$^{+}$ cells magnetically isolated from the bone marrow of P1/2 KO versus P1/2-CTR mice**, related to Figures 5, 6, and S6.
- Table S5: **Enrichment analysis of proteomics data**, related to Figures 5, 6 and S6.
- Table S6: **Panels of antibodies and dyes used in flow cytometry analysis.**

***STAR*METHODS**

**KEY RESOURCES TABLE**

| REAGENT or RESOURCE | SOURCE | IDENTIFIER |
|---|---|---|
| **Antibodies** | | |
| Beta ACTIN Monoclonal Antibody (Clone AC-15) | Sigma-Aldrich | Cat# A1978; RRID: AB_476692 |
| CD4 Monoclonal Antibody (clone GK1.5), PE | Thermo Fisher | Cat# 12-0041-81; RRID: AB_465505 |
| CD4 Monoclonal Antibody (clone GK1.5), PE-Cy7 | Thermo Fisher | Cat# 25-0041-82; RRID: AB_469576 |
| CD8a Monoclonal Antibody (clone 53-6.7), PE | Thermo Fisher | Cat# 12-0081-81; RRID: AB_465529 |
| CD8a Monoclonal Antibody (clone 53-6.7), PE-Cy7 | Thermo Fisher | Cat# 25-0081-81; RRID: AB_469583 |
| CD9 Monoclonal Antibody (clone KMC8), FITC | Thermo Fisher | Cat# 11-0091-82; RRID: AB_1210669 |
| CD11b Monoclonal Antibody (Clone M1/70), APC | Thermo Fisher | Cat# 17-0112-81; RRID: AB_469342 |
| CD11b Monoclonal Antibody (Clone M1/70), PE-Cy7 | Thermo Fisher | Cat# 25-0112-81; RRID: AB_469587 |
| CD11b Monoclonal Antibody (Clone M1/70), Pacific Blue | BioLegend | Cat# 101224; RRID: AB_755986 |
| CD11c Monoclonal Antibody (Clone N418), FITC | BioLegend | Cat# 117305; RRID: AB_313774 |
| CD127 Monoclonal Antibody (Clone A7R34), PE | Thermo Fisher | Cat# 12-1271-81; RRID: AB_465843 |
| CD16/32 Monoclonal Antibody (Clone 93), eFluor 450 | Thermo Fisher | Cat# 48-0161-82; RRID: AB_1272191 |
| CD16/32 Monoclonal Antibody (Clone 93), PE | Thermo Fisher | Cat# 12-0161-82; RRID: AB_465568 |
| CD34 Monoclonal Antibody (Clone RAM34), FITC | Thermo Fisher | Cat# 11-0341-85; RRID: AB_465022 |
| CD34 Monoclonal Antibody (Clone RAM34), Alexa Fluor 700 | Thermo Fisher | Cat# 56-0341-82; RRID: AB_493998 |
| CD41a Monoclonal Antibody (eBioMWReg30), eFluor 450 | Thermo Fisher | Cat# 14-0411-82; RRID: AB_ AB_763490 |
| CD42d Monoclonal Antibody (Clone 1C2), APC | Thermo Fisher | Cat# 17-0421-82; RRID: AB_1724071 |
| CD44 Monoclonal Antibody (Clone IM7), PE | Thermo Fisher | Cat# 12-0441-82; RRID: AB_465664 |
| CD45 Monoclonal Antibody (Clone 30-F11), Pacific Blue | BioLegend | Cat# 103126; RRID: AB_493535 |
| CD45.1 Monoclonal Antibody (Clone A20), Pacific Blue | BioLegend | Cat# 110722; RRID: AB_492866 |
| CD45.2 Monoclonal Antibody (Clone 104), FITC | Thermo Fisher | Cat# 11-0454-81; RRID: AB_465060 |
| CD45R (B220) Monoclonal Antibody (Clone RA3-6B2), PE | Thermo Fisher | Cat# 12-0452-81; RRID: AB_465670 |
| CD45R (B220) Monoclonal Antibody (Clone RA3-6B2), APC | Thermo Fisher | Cat# 17-0452-81; RRID: AB_469394 |
| CD45R (B220) Monoclonal Antibody (Clone RA3-6B2), PE-Cy7 | Thermo Fisher | Cat# 25-0452-81; RRID: AB_469626 |
| CD48 Monoclonal Antibody (Clone HM48-1), PE | Thermo Fisher | Cat# 12-0481-81; RRID: AB_465693 |
| CD48 Monoclonal Antibody (Clone HM48-1), Pacific Blue | BioLegend | Cat# 103417; RRID: AB_756139 |
| CD71 (TFRC) Monoclonal Antibody (Clone RI7217), FITC | BioLegend | Cat# 113806; RRID: AB_313567 |
| CD117 (c-KIT) Monoclonal Antibody (Clone 2B8), PE | BioLegend | Cat# 105808; RRID: AB_313217 |
| CD117 (c-KIT) Monoclonal Antibody (Clone 2B8), APC | Thermo Fisher | Cat# 17-1171-81; RRID: AB_469429 |
| CD135 Monoclonal Antibody (Clone A2F10), PE | Thermo Fisher | Cat# 12-1351-82; RRID: AB_465859 |
| CD150 Monoclonal Antibody (Clone TC15-12F12.2), PE-Cy5 | BioLegend | Cat# 115911; RRID: AB_493599 |
| CD170 Monoclonal Antibody (Clone S17007L), PE | BioLegend | Cat# 155505; RRID: AB_2750234 |
| F4/80 Monoclonal Antibody (Clone BM8), Pacific Blue | BioLegend | Cat# 123124; RRID: AB_893475 |
| FcεRIα Monoclonal Antibody (Clone MAR-1), Pacific Blue | BioLegend | Cat# 134313; RRID: AB_10612933 |
| Ferritin Heavy Chain Monoclonal antibody | Abcam | Cat# ab183781 |
| Ferritin Light Chain Polyclonal antibody (Clone EPR18878) | Abcam | Cat# ab69090; RRID: AB_1523609 |
| Ferroportin Polyclonal Antibody | Alpha Diagnostics | Cat# MTP11-A; RRID: AB_1619475 |
| IA/IE (MHCII) Monoclonal Antibody (Clone M5/114.15.2), PE | Biolegend | Cat# 107607; RRID: AB_313322 |
| LY6A/E (SCA1) Monoclonal Antibody (Clone D7), APC-Cy7 | BD Biosciences | Cat# 560654; RRID: AB_1727552 |
| LY6C Monoclonal Antibody (Clone HK1.4), PE | BioLegend | Cat# 128007; RRID: AB_1186133 |
| LY6C Monoclonal Antibody (Clone HK1.4), PE-Cy7 | BioLegend | Cat# 128018; RRID: AB_1732082 |
| LY6G (Gr-1) Monoclonal Antibody (Clone RB6-8C5), PE-Cy7 | Thermo Fisher | Cat# 25-5931-81; RRID: AB_469662 |

| | | |
|---|---|---|
| LY6G Monoclonal Antibody (Clone 1A8), Alexa Fluor488 | BioLegend | Cat# 127626; RRID: AB_2561340 |
| LY6G Monoclonal Antibody (Clone 1A8), APC-Cy7 | BioLegend | Cat# 127624; RRID: AB_10640819 |
| TER119 Monoclonal Antibody (Clone TER-119), PE-Cy7 | Thermo Fisher | Cat# 25-5921-82; RRID: AB_469661 |
| Chemicals, peptides, and recombinant proteins | | |
| Neupogen 300 µg/0.5 mL | Amgen | Cat# 1011921 |
| Sunflower seed oil | Sigma-Aldrich | Cat# 88921 |
| Tamoxifen free base | Sigma-Aldrich | Cat# T5648 |
| Ferriprox | Apotex Europe B.V | N/A |
| Ferric Ammonium Citrate | Sigma-Aldrich | Cat# F5879 |
| Deferiprone | Sigma-Aldrich | Cat# 379409 |
| Recombinant Murine SCF (KITL) | PeproTech | Cat# 250-03 |
| Recombinant Murine IL3 | PeproTech | Cat# 213-13 |
| Recombinant Murine M-CSF (CSF1) | PeproTech | Cat# 315-02 |
| Recombinant Murine GM-CSF (CSF2) | PeproTech | Cat# 315-03 |
| Recombinant Murine G-CSF (CSF3) | PeproTech | Cat# 250-05 |
| 2-NBDG | PeproTech | Cat# 1860768 |
| BioTracker™ Far-red Labile Fe2+ Dye | Merck Millipore | Cat# SCT037 |
| BODIPY 493/503 | Thermo Fisher | Cat# D3922 |
| CellROX Green Reagent | Thermo Fisher | Cat# C10444 |
| ACK lysing buffer | Biozym | Cat# 882090 |
| Cytochalasin D | Santa Cruz | Cat# sc-201442 |
| JC-1 | Enzo Life Sciences | Cat# ENZ-52304 |
| Lipopolysaccharide (LPS) | Sigma-Aldrich | Cat# L4641 |
| MitoTracker™ Green FM | Thermo Fisher | Cat# M7514 |
| pHrodo Green S. Aureus Bioparticles | Thermo Fisher | Cat# P35367 |
| Thiazol Orange | Sigma-Aldrich | Cat# 390062 |
| Zombie UV™ Fixable Viability | BioLegend | Cat# 423108 |
| FCCP | Sigma-Aldrich | Cat# C2920 |
| Foetal Bovine Serum (low endotoxin) | VWR International | Cat# S1860 |
| IMDM | LIFE Technologies | Cat# 21980032 |
| L-Glutamine | VWR International | Cat# K0282 |
| Penicillin-Streptomycin | Thermo Fisher | Cat# 15140122 |
| DAPI | Thermo Fisher | Cat# D1306 |
| iTaq Universal SYBR green supermix | BIO-RAD | Cat# 1725125 |
| Halt™ Protease Inhibitor Cocktail | Thermo Fisher | Cat# 87786 |
| Halt™ Phosphatase Inhibitor Cocktail | Thermo Fisher | Cat# 78427 |
| Critical commercial assays | | |
| Mouse Erythropoietin/EPO Quantikine ELISA Kit | Bio-techne | Cat# MEP00B |
| Hepcidin Murine-Compete™ ELISA kit | Intrinsic LifeSciences | Cat# HMC-001 |
| Mouse Haptoglobin ELISA Kit | Abcam | Cat# ab157714 |
| Mouse Hemopexin ELISA Kit | Abcam | Cat# ab157716 |
| Extracta™ DNA Prep for PCR - Tissue | VWR International | Cat# 733-2160 |
| Arcturus®PicoPure® RNA isolation kit | Thermo Fisher | Cat# KIT0204 |
| RNase-Free DNase Set | Qiagen | Cat# 79254 |
| RNA 6000 pico kit | Agilent | Cat# 5067-1513 |
| SuperScript™ VILO™ cDNA Synthesis Kit | Thermo Fisher | Cat# 11754050 |
| FITC BrdU Flow Kit | BD Biosciences | Cat# 559619; RRID: AB_2617060 |
| CYTO-ID® Autophagy detection kit | Enzo Life Sciences | Cat# ENZ-51031-0050 |
| Direct Lineage Cell Depletion Kit | Miltenyi Biotec | Cat# 130-110-470 |
| Anti-Ly-6G microBeads ultra pure | Miltenyi Biotec | Cat# 130-120-337 |
| FASTSTART PCR Master mix | Sigma-Aldrich | Cat# 4710452001 |
| SMART-Seq® v4 Ultra® Low Input RNA Kit | Takara | Cat# R400752 |

| | | |
|---|---|---|
| **Deposited data** | | |
| RNA-seq fastq data | This paper | E-MTAB-10663<br>www.ebi.ac.uk/arrayexpress |
| Proteomics data | This paper | PXD032077<br>http://www.ebi.ac.uk/pride |
| Granule protein annotation | Grassi et al., 2018 | Table S6<br>DOI: 10.1016/j.celrep.2018.08.018 |
| Mitocarta 3.0 database | Rath et al., 2021 | DOI: 10.1093/nar/gkaa1011 |
| Proteomics data during neutropoiesis in humans | Hoogendijk et al., 2019 | Table S1<br>DOI: 10.1016/j.celrep.2019.10.082 |
| Mouse reference genome GRCm38.p5 | Genome Reference Consortium | www.ensembl.org |
| Ribosomal protein gene database | http://ribosome.med.miyazaki-u.ac.jp/ | Cytoplasmic Ribosomal Proteins (Mus Musculus) |
| **Experimental models: Organisms/strains** | | |
| Mouse; *Aco1*/*Irp1* flox; B6.Cg-*Aco1*$^{tm1.1Mwh}$ | Galy et al., 2005a | MGI:3613495 |
| Mouse; *Ireb2*/*Irp2* flox; B6.Cg-*Ireb2*$^{tm1.1Mwh}$ | Galy et al., 2005a | MGI:3613523 |
| Mouse; Rosa26 CreERT2; B6.Cg-Gt(ROSA)26$^{Sortm1(cre/ERT)Nat/J}$ | Jackson Laboratories | JAX: #004847 |
| Mouse; wild type C57BL6/J (used in Ferriprox experiments) | JANVIER LABS | C57BL/6JRj |
| Mouse; Ly5.1, B6 Cd45.1, B6.SJL-Ptprc$^{a}$ Pepc$^{b}$/BoyJ (CD45.1$^{+}$ mice used in transplantation experiments) | Jackson Laboratories | JAX #002014 |
| **Oligonucleotides** | | |
| *Aco1* genomic PCR-forward 1<br>5’-TACTGTAGCAAAAATGCTTTGTCTCTG-3’ | Sigma Oligo | N/A |
| *Aco1* genomic PCR-forward 2<br>5’-GTCATTTTTCTCATTTCTTGAGCATTAG-3’ | Sigma Oligo | N/A |
| *Aco1* genomic PCR-reverse<br>5’-TCTATCCCTGAGGTCGGTAGGC-3’ | Sigma Oligo | N/A |
| *Ireb2* genomic PCR-forward 1<br>5’-CTGAAAGACTGACCCTTCTTGTTC-3’ | Sigma Oligo | N/A |
| *Ireb2* genomic PCR-forward 2<br>5’-TGAGTGGTGCCTGCATTTTAAG-3’ | Sigma Oligo | N/A |
| *Ireb2* genomic PCR-reverse<br>5’-GGCTTCAATAGTCTTCATACCACG-3’ | Sigma Oligo | N/A |
| *Gusb* qRT-PCR-forward<br>5’-AAAATCACCCTGCGGTTGT-3’ | Sigma Oligo | N/A |
| *Gusb* qRT-PCR- reverse<br>5’-TGTGGGTGATCAGCGTCTT-3‘ | Sigma Oligo | N/A |
| *Ppib*-qRT-PCR- forward<br>1 5’-GGAGATGGCACAGGAGGA-3‘ | Sigma Oligo | N/A |
| *Ppib* qRT-PCR-reverse<br>5‘-GGTGTCTTTGCCTGCATTG-3‘ | Sigma Oligo | N/A |
| PrimePCR™ SYBR® Green Assay: *Ubxn4* | BIO-RAD | qMmuCID0023115 |
| PrimePCR™ SYBR® Green Assay: *Psmc2* | BIO-RAD | qMmuCIP0034703 |
| PrimePCR™ SYBR® Green Assay: *Abcg1* | BIO-RAD | qMmuCID0005670 |
| PrimePCR™ SYBR® Green Assay: *Bhlhe40* | BIO-RAD | qMmuCID0013865 |
| PrimePCR™ SYBR® Green Assay: *Clec7a* | BIO-RAD | qMmuCID0013926 |
| PrimePCR™ SYBR® Green Assay: *Egln3* | BIO-RAD | qMmuCID0019903 |
| PrimePCR™ SYBR® Green Assay: *Mylip* | BIO-RAD | qMmuCID0010376 |
| PrimePCR™ SYBR® Green Assay: *Pfkl* | BIO-RAD | qMmuCID0018472 |
| PrimePCR™ SYBR® Green Assay: *Pfkp* | BIO-RAD | qMmuCID0023075 |
| PrimePCR™ SYBR® Green Assay: *Slc2a1* | BIO-RAD | qMmuCED0026836 |
| **Software and algorithms** | | |
| FlowJo v.10 | Tree Star | www.flowjo.com |

| | | |
|---|---|---|
| GraphPad Prism V9 | www.graphpad.com | N/A |
| STAR Package | Dobibn et al., 2013 | http://code.google.com/p/rna-star/. |
| htseq-count package | Anders et al., 2015 | https://pypi.org/project/HTSeq/ |
| Limma software | Ritchie et al., 2015 | http://bioinf.wehi.edu.au/limma |
| missForest package | Stekhoven anmd Bühlmann, 2012 | https://github.com/stekhoven/missForest |
| FastQC | Andrews S (2010) | www.bioinformatics.babraham.ac.uk/projects/fastqc/ |
| DESeq2 package | Love et al., 2014 | www.bioconductor.org/packages/release/bioc/html/DESeq2.html |
| topGO package | Alexa and Rahnenfuhrer | DOI: 10.18129/B9.bioc.topGO |
| ggplot2 package | Bioconductor | https://CRAN.R-project.org/package=ggplot2 |
| MaxQuant v 1.6.14.0 | Tyanova et al., 2016a | https://www.maxquant.org/maxquant/ |
| Perseus software | Tyanova et al., 2016b | https://www.maxquant.org/perseus/ |

### LEAD CONTACT AND MATERIALS AVAILABILITY

This study did not produce new unique reagents.

### DATA AND CODE AVAILABILITY

RNA-seq data have been deposited in the ArrayExpress database at EMBL-EBI (www.ebi.ac.uk/arrayexpress) under accession number E-MTAB-10663.
The mass spectrometry proteomics data have been deposited to the ProteomeXchange Consortium via the PRIDE (Perez-Riverol et al., 2022) partner repository (http://www.ebi.ac.uk/pride) with the dataset identifier PXD032077.

### EXPERIMENTAL MODEL AND SUBJECT DETAILS

#### Mice

*Husbandry*:
The mouse lines carrying floxed Aco1 ($Aco1^{tm1.1Mwh}$) and Ireb2 ($Ireb2^{tm1.1Mwh}$) alleles and the $Gt(ROSA)26Sor^{tm1(cre/ERT)Nat}$ deletor strain expressing a tamoxifen-inducible CRE recombinase (CreER) under the control of the *Rosa26* promoter have been described (Galy et al., 2005a, Badea et al., 2003). $\mathit{Aco1}^{flox/flox}\mathit{Ireb2}^{flox/flox}\mathit{Rosa26}^{+/CreERT2}$ (P1/2-KO) and $\mathit{Aco1}^{flox/flox}\mathit{Ireb2}^{flox/flox}\mathit{Rosa26}^{+/+}$ (P1/2-CTR) littermates were used throughout the study. To control for potential effects of CRE, $\mathit{Aco1}^{+/+}\mathit{Ireb2}^{+/+}\mathit{Rosa26}^{+/CreERT2}$ (CreER) and wild type $\mathit{Aco1}^{+/+}\mathit{Ireb2}^{+/+}\mathit{Rosa26}^{+/+}$ (WT) littermates were also analyzed. All mice were on a homogenous C57BL6/J genetic background and were housed under specific pathogen-free and light-, temperature- (21°C), and humidity (50-60% relative humidity)-controlled conditions. Food (containing around 200 ppm of iron) and water were available *ad libitum*. Blood was collected by cardiac puncture after deep anesthesia of the mice with a Ketamin/Xylazin mixture. If not used immediately, tissues were flash-frozen in liquid nitrogen and stored at -80°C until analysis.

*Treatments*:
To induce CRE activity during adulthood, 8-12 week-old male mice were injected intraperitoneally with a low dose of tamoxifen (40 mg/kg body weight in a 1:9 EtOH-sunflower seed oil mix, all from Sigma-Aldrich Chemie GmbH, Munich, Germany) on day 1 and day 3. Compared to the standard regimen of daily injections of 175 mg/kg/day on 5 consecutive days (Higashi et al., 2009), we found that this milder treatment triggers efficient recombination of floxed substrates while minimizing CRE toxicity (unpublished data). To decrease body iron availability in adult animals, 8 week-old C57BL6/J wild type mice received deferiprone (Ferriprox™ from Apotex Europe B.V., Leiden, Holland) in the drinking water at a dose of 0.4 µg/kg body weight per day for a period of 2 months. They were subsequently treated daily with G-CSF (Neupogen® from Amgen GmbH, Munich, Germany) at a dose of 250 µg/kg body weight on 3 consecutive days in order to stimulate granulopoiesis; the animals were sacrificed one day after the last G-CSF injection for analysis. To label proliferating cells, the mice were injected (i.p.) with BrdU (BD Biosciences, Heidelberg, Germany) at a dose of 50 µg/kg body weight 2 hours prior sacrifice.
*Bone marrow transplantation and generation of chimeric mice:*
The mice were exposed to a total of 10 Gy (in two doses separated by a 3 to 4 hours interval) from a $^{137}Cs$ source. Following a 2 hours recovery period, the mice were injected (i.v.) with $6x10^6$ total BM cells from donor animals. As a prophylactic measure, the transplanted animals were treated with carprofen (s.c., 5 mg/kg body weight) once a day for the first 3 days and received sulfamethoxazole/trimethoprim (90 mg/kg/day) in the drinking water for 3 weeks. They were additionally given soft food. To assess engraftment efficiency, small blood samples were collected from the submandibular vein into EDTA tubes (Sarstedt, Nümbrecht, Germany) once a month, and the samples were analyzed by flow cytometry as described below. Successful engraftment was typically obtained 12 weeks after transplantation.
*Ethical statement:*
Animal care, husbandry, and killing were performed according to national guidelines and were approved by an institutional review board headed by the local animal welfare officers. Animal experiments were carried out according to project licenses G-270/18 and G-21/21, as approved by the Regiergrungspräsidium of Karlsruhe (Baden-Württemberg, Germany).

### METHOD DETAILS

#### Hematology and serum parameters
Blood profiles and hemoglobin content were determined using an ABC Vet apparatus (HORIBA ABX SAS, Montpellier, France). Serum samples were prepared using Z-gel containing microvette® tubes (Sarstedt, Nümbrecht, Germany). EPO concentration was determined with the Mouse Erythropoietin Quantikine ELISA kit (Bio-Techne, Wiesbaden, Germany). Hepcidin was measured using a Hepcidin Murine-Compete™ ELISA kit (Intrinsic LifeSciences, La Jolla, CA). Haptoglobin and Hemopxin levels were analyzed using, respectively, the Mouse Haptoglobin ELISA (ab157714) and Mouse Hemopexin (ab157716) ELISA kits from Abcam (Cambridge, UK). Serum concentration of iron, ferritin and transferrin was determined at the "Centre de Recherche sur l'Inflammation" (Paris, France) using an Olympus 400 analyzer (Olympus, Tokyo, Japan).

#### Tissue iron levels
Spleen and liver tissues were dried and non-heme iron levels were determined against a standard using the bathophenanthroline chromogen as described previously (Tybl et al., 2020).

#### DNA analysis
DNA was isolated using the Extracta DNA Prep kit from Quantabio (VWR International, Darmstadt, Germany). The genomic status of the *Aco1* and *Ireb2* loci was analyzed by PCR using PCR Mastermix reagents (Merck KGaA, Darmstadt, Germany) together with the primers listed in Key Resources Table. The amplicons obtained have the following size: wild type *Aco1* allele; 243 bp; floxed *Aco1* allele, 296 bp; truncated *Aco1* allele, 214 bp; wild type *Ireb2* allele, 245 bp; floxed *Ireb2* allele, 288 bp; truncated *Ireb2* allele, 199 bp.

**RNA analyses**

*RNA extraction.*

Total RNA from sorted Prog, prNeu and PMN cell populations was extracted using the Arcturus®PicoPure® RNA isolation kit (Thermo Fisher Scientific, Darmstadt, Germany) and treated with RNAse-free DNAseI (Qiagen, Düsseldorf, Germany) to eliminate genomic DNA. RNA concentration was measured with a Qubit Assay (Thermo Fisher Scientific) and RNA integrity was assessed using an RNA 6000 pico kit together with a Bioanalyser apparatus from Agilent (Agilent Technologies, Waldbronn, Germany).

*qRT-PCR.*

Total RNA was reverse-transcribed using the SuperScript VILO Master kit (Thermo Fisher Scientific) and RNA levels were determined on a CFX Connect Real-Time PCR System using the iTaq Universal SYBR Green Supermix (Bio-Rad Laboratories GmbH, Feldkirchen, Germany) together with the primers listed in Key Resources Table and the following cycling parameters: denaturation at 95°C for 10 min, followed by 45 cycles of 15 s at 95°C, 15 s at 60°C and 15 s at 72° C. For each assay, primer efficiency (E) was determined using a serial dilution of a mix of cDNA from a pool of all samples analyzed. Semi-quantification of a gene of interest (GOI) was done using the ΔΔCt-method after calibration to the average expression of four standard genes (Std) as indicated in figure legends. The expression fold change was calculated as: $[E_{GOI}*(\Delta Ct_{GOI})]/[E_{Std}*(\Delta Ct_{Std})]$.

*RNA sequencing analysis:*

Ultra-Low RNA-seq libraries were prepared from 2.85 ng input RNA using the SMART-Seq® v4 Ultra® Low Input RNA Kit for Sequencing (Takara Bio Europe SAS, Saint-Germain-en-Laye, France), combined with shearing on a Covaris apparatus (Covaris Ltd., Brighton, UK) and the NEBNext® ChIP-Seq Library Prep Master Mix Set for Illumina (New England Biolabs GmbH, Frankfurt am Main, Germany) according to the manufacturer's protocol. The libraries were quality controlled using an Agilent 4200 Tape Station System (Agilent Technologies) and a Qubit ds DNA HS Assay kit (Thermo Fisher Scientific). Based on Qubit quantification and sizing analysis, libraries were normalized, pooled, and clustered on a cBot system (Illumina) with a final concentration of 250 pM (spiked with 1% PhiX control v3 from Illumina). 50 bp single-read sequencing was performed on a HiSeq 4000 instrument (Illumina) using standard protocols.

Demultiplexed Fastq files were checked for quality with the FastQC program (Andrews, 2010), and mapped against the mm10 mouse genome using the STAR (Spliced Transcripts Alignment to a Reference) package (Dobin et al., 2013) to generate bam files. Sequencing reads were annotated against the GRCm38.p5 reference gene set from Ensembl (www.ensembl.org) and counted using the htseq-count package (Anders et al. 2015). Differentially expressed genes were identified with the DESeq2 package (Love at al., 2014), using the Benjamini-Hochberg (BH) adjustment for false discovery rate (FDR) calculation (Benjamini and Hochberg, 1995). Principal Component Analysis (PCA) plot was generated using the plotPCA function of the DESeq2 package.

RNA-seq fastq files can be obtained from the Arrayexpress platform (www.ebi.ac.uk/arrayexpress/) under the accession number E-MTAB-10663.

**Western blotting**

Magnetically sorted $LIN^-$ versus $LY6G^+$ cells were lysed in a RIPA buffer (50 mM tris-HCl pH7.5, 150 mM NaCl, 1% Triton X-100, 0.5% Na deoxycholate, 0.1% SDS, 1mM DTT) supplemented with the Halt™ Protease and Halt™ Phosphatase Inhibitor Cocktails (Thermo Fisher Scientific). The samples were incubated on ice for 30 min with occasional vortexing and the debris was pelleted at 10000 *g* for 10 min at 4°C. Protein concentration in the supernatant was determined using the Pierce™ BCA protein Assay kit (Thermo Fisher Scientific). Equal amounts of protein were mixed with Laemmli sample buffer, resolved onto 4-15% Criterion™ TGX™ midi protein gels, and transferred onto PVDF membranes using a Trans-Blot® Turbo™ 0.2 µm PVDF transfer pack together with a Trans-Blot Turbo transfer system (all from Bio-Rad laboratories). The membranes were incubated in TBS-T (Tris-buffered saline, 0.1% Tween-20) containing 5% (w/v) powder milk and the antibodies listed in Key Resources Table. Immune complexes were visualized using the Clarity™ western ECL substrate (Bio-Rad laboratories) together with a Chemocam Imager system equipped with the ChemoStar Imager software (INTAS Science Imaging Instruments GmbH, Göttingen, Germany).

**Proteomics analysis**

*Sample preparation and mass spectrometry (MS).*

An amount of 10 ug of protein per condition was usef for a short SDS-PAGE separation (0.5 cm). After Commassie staining, the total protein sample was excised and subjected to trypsin digestion on a DigestPro MSi robotic system (CEM GmbH, Kamp-Lintfort, Germany) as described previously (Shevchenko et al., 2006). The LC-MS/MS analysis was performed on an Ultimate 3000 UPLC system directly connected to an Orbitrap Exploris 480 mass spectrometer (both from Thermo Fisher Scientific). Peptides were desalted on an Acclaim PepMap300 C18 trap cartridge (5µm, 300Å wide pore from Thermo Fisher Scientific) for 3 min using a 30 µL/min flow of 0.05% trifluoroacetic acid in water. The analytical multistep gradient was carried out on a nanoEase MZ Peptide analytical column (300Å, 1.7 µm, 75 µm x 200 mm) from Waters GmbH (Eschborn, Germany) using 0.1% formic acid in water as solvent A and 0.1% formic acid, 80% acetonitrile in water as solvent B. Solvent B concentration was linearly increased from 2% to 38% during 134 min, followed by a quick raise to 95% for two minutes. Solvent B concentration was then lowered to 2% and a 10 min equilibration step was added. Eluting peptides were analyzed in the mass spectrometer using a data depend acquisition (DDA) mode. A full scan at 60k resolution (380-1400 m/z, 300% AGC target, 45 ms maxIT) was followed by up to 2 seconds of MS/MS scans. Peptide features were isolated with a window of 1.4 m/z, fragmented using 26% normalized collision energy. Fragment spectra were recorded at 15k resolution (100% automatic gain control target, 54 ms maxIT). Unassigned and singly charged eluting features were excluded from fragmentation and dynamic exclusion was set to 35s.

*Data analysis*.

Data analysis was carried out by MaxQuant (version 1.6.14.0, Tyanova et al., 2016a) using an organism specific database extracted from Uniprot.org under default settings. Identification FDR cutoffs were 0.01 on peptide level and 0.01 on protein level. Match between runs option was enabled but restricted to transfer peptide identifications across RAW files based on accurate retention time and m/z only within replicates.
LFQ quantification was done using a label free quantification approach based on the MaxLFQ algorithm (Cox et al, 2014). A minimum of 2 quantified peptides per protein was required for protein quantification.

**Flow cytometry (FCM)**

The samples were analyzed on an LSR Fortessa device (BD Biosciences) and the data processed using the Flow-Jo software v.10 (Tree Star, Ashland, OR).

*Cell preparation and surface marker staining:*

To analyze BM cells, the femur, tibiae and ilia were disinfected with 70% EtOH and crushed in Iscove Modified Dulbecco Medium (Thermo Fisher Scientific) using a mortar and pestle. The BM cell suspension obtained was filtered through a 40 µm cell strainer (Greiner Bio-one, Frickenhausen, Germany) and nucleated cells were counted using a Hemavet apparatus (Drew Scientific, Miami Lakes, FL). Following a PBS wash, cells were stained for 20 min at RT with a zombie dye (BioLegend, Koblenz, Germany) to discriminate live versus dead cells. For surface marker staining, cells were washed in PBS and subsequently incubated for 20 min at 4°C in PBS containing 2% fetal bovine serum (FBS, Thermo Fischer Scientific) and antibody mixes as listed in table S6 (panels P1 to P21). To get rid of red cells prior analysis, the samples were incubated in ACK red cell lysis buffer (Biozym Scientific GmbH, Hessisch Oldendorf, Germany) for 10 min at RT and then washed in PBS. For *ex vivo* differentiation experiments (see below), the cells were washed and stained as described above, using the antibody panels P22 and P23 (Table S6); DAPI was added to the cells shortly before FCM.

*Reticulocyte counts in PB*:

PB cells were washed and resuspended in PBS + 2% FBS containing anti-CD45 and anti-TER119 antibodies (panel P8, Table S6), and were incubated for 30 min on ice. Following a PBS wash, the cells were resuspended in PBS 2%FBS complemented with 1.7 µg/mL thiazol orange (Sigma-Aldrich) to stain the residual RNA present in reticulocytes. After a 30 min incubation on ice, the cells were directly analyzed by FCM.

*Assessment of BM cell engraftment*:
PB cells (collected from the submandibular vein) were incubated for 20 min on ice in PBS + 2%FBS containing fluorochrome-conjugated antibodies against leukocyte antigens and the CD45.1/CD45.2 polymorphism markers (panels P5 and P6, Table S6). Prior analysis, red blood cells were eliminated using the ACK buffer as described above.

*Cell cycle analysis based on BrdU incorporation:*
BM cells were incubated with antibodies against cell surface markers (panel P16, Table S6), and were additionally stained for BrdU using the FITC BrdU Flow kit (BD Biosciences) according to the manufacturer's instructions. Briefly, cells were washed in PBS + 2%FCS and fixed/permeabilized in cytofix/cytoperm solution, washed with Perm/Wash, and incubated with the anti-BrdU antibody overnight at 4°C. DNase treatment was used to expose the BrdU epitope. To determine the DNA content, BrdU-stained cells were incubated with DAPI (4' 6-Diamidino-2-Phenylindole, Thermo Fisher Scientific).

*Study of the "labile iron pool":*
BM cells washed in PBS were incubated for 1h at 37°C in serum-free IMDM supplemented with 5 µM of BioTracker™ Far-red Labile ferrous iron Dye (Merck KGaA). Cells were then washed in PBS and stained for cell surface markers (panel P14, Table S6) as described above.

*Analysis of neutral lipid content:*
BM cells washed in PBS were incubated 30 min at 37°C in IMDM containing 1 µM of the fluorescent neutral lipid dye BODIPY 493/503 (4,4-difluoro-1,3,5,7,8-pentamethyl-4-bora-3a,4a-diaza-s-indacene) (Thermo Fischer Scientific). Cells were then washed in PBS and stained for cell surface markers (panel P15, Table S6) as described above.

*Autophagy detection:*
$LIN^-$ and $LY6G^+$ cells were magnetically isolated as described below. The sorted cells were washed and then incubated for 30 min at 37°C in assay buffer containing 1:1000 of the CYTO-ID dye (Enzo Life Sciences GmbH, Lörrach, Germany), a cationic amphiphilic tracer that labels autophagic vacuoles. The cells were analyzed by FCM after a PBS wash.

*Monitoring of 2-NBDG glucose uptake:*
BM cells washed in PBS were incubated for 1 h at 37°C in glucose-free RPMI containing 100 µM of 2-NBDG (2-deoxy-2-[(7-nitro-2,1,3-benzoxadiazol-4-yl)amino]-D-glucose) (Thermo Fischer Scientific). Cells were then washed in PBS and stained for cell surface markers (panel P17, Table S6) as described above. ROS detection was validated using cells treated with tert-butyl hydroperoxide as positive control.

*Analysis of ROS production*:
BM cells in serum-free IMDM were stimulated with lipopolysaccharide (LPS) at a concentration of 2.5 µg/mL. After a 1 h incubation at 37°C, the cells were washed in PBS, pelleted at 350 *g* for 5 min then resuspended in serum-free IMDM containing 5 µM CellROX™ green reagent (ThermoFisher scientific). The cells were then incubated for 20 min at 37°C. Cells were washed in PBS and stained for cell surface markers (panel 20, Table S6) as described above.

*Study of phagocytic activity*:
BM cells were incubated for 1h at 37°C in serum-free IMDM in the presence of LPS (2.5 µg/mL) and pHrodo™ Green S. aureus Bioparticles™ Conjugate (ThermoFisher scientific) at a 1:10 ratio. The cells were then washed in PBS, centrifuged at 350 *g* for 5 min, washed again in PBS and stained for cell surface markers (panel 21, Table S6) as described above. As controls, the assay was performed either in the presence of 10 µM of cytochalasin D (sc-201442, Santa Cruz Biotechnology Inc., Heidelberg, Germany) to block F-actin polymerization and phagocytosis, or with cells incubated on ice.

*Analysis of Mitochondrial status:*
BM cells were washed in PBS and pelleted. Cells were then incubated at 37°C for 45 min in IMDM containing 200 nM of Mitotracker Green dye (Thermo Fisher scientific) to assess mitochondrial mass. To analyze the mitochondrial membrane potential, the cells were incubated at 37°C for 15 min in IMDM supplemented with JC-1 reagent at a concentration of 10 µg/mL. After a PBS wash, the cells were stained for surface markers (panels P18 and P19, Table S6) as described above. To control for JC-1 red fluorescence specificity, cells were pre-treated for 45 min with 250 nM of the depolarizing agent FCCP (Carbonyl cyanide-p-trifluoromethoxyphenylhydrazone, C2920, Sigma-Aldrich).

**Cell sorting**
*Fluorescence-activated cell sorting (FACS):*
BM cells were prepared and stained with antibodies against cell surface markers as described above for standard FCM. The cell populations of interest were isolated using a FACSAria™ Cell Sorter (BD). A fraction of the sorted cells was re-analyzed by FCM to ascertain the purity (typically > 90%) of the cell populations obtained. For DNA and RNA extraction, respectively, the sorted cells were collected directly into the corresponding lysis buffers. For imaging, the cells were collected in PBS + 5%FBS.

*Magnetic sorting:*
BM cell suspensions were prepared as described for FCM analysis. Hematopoietic progenitor cells were isolated using the Direct lineage depletion kit for mouse (Miltenyi Biotec, Bergisch Gladbach, Germany). $2x10^7$ BM cells in 80µL of MACS buffer (PBS + 0.5% FBS + 2 mM EDTA) were incubated for 10 minutes at 4°C with 20 µL of microbeads conjugated to antibodies against various differentiation markers. The labelled cells were then loaded onto LS columns (Miltenyi Biotec) pre-equilibrated with MACS buffer and placed on a QuadroMACS™ Separator (Miltenyi Biotec). The flowthrough containing progenitor cells was collected for downstream applications.
To isolate $LY6G^+$ cells, $10^7$ BM cells in 90 µL of MACS buffer were incubated for 10 min at 4°C with 10 µL of Anti-LY6G MicroBeads UltraPure for mouse (Miltenyi Biotech). The cells were then loaded onto LS columns to eliminate the $LY6G^-$ cells present in the flowthrough. The columns were subsequently washed with MACS buffer, and $LY6G^+$ cells were eluted in the same buffer.

**Cell culture**
BM progenitor cells were isolated using the Direct lineage depletion kit as described earlier. The cells were seeded at a density of $2x10^5$ cells per mL in progenitor growth medium (PGM) containing IMDM (Thermo Fisher Scientific), 10% heat-inactivated and low endotoxin FBS (VWR International), 1% Penicillin/Streptomycin (P/S, Thermo Fisher Scientific), and 5% L-Glutamine (VWR International). To enable the maintenance and expansion of progenitor cells, the culture medium was supplemented with interleukin 3 (IL3) and stem cell factor (SCF, a.k.a. KITL). Cells were incubated at 37°C in 5% $CO_2$. Following 3 days of expansion, the cells were cryopreserved in 90%FBS + 10% dimethyl sulfoxide for later use.
To study cell differentiation, progenitor cells were thawed in PGM containing IL3 + SCF and cultivated for 2 to 3 days in 6-well plates. After this recovery phase, the cells were seeded in 12-well plates and grown in the same medium for another 3 days. To initiate neutrophil differentiation, the cells were washed in PBS and cultivated for 2 days in PGM containing IL3+SCF and complemented with G-CSF (granulocyte colony stimulating factor, a.k.a. CSF3). For the final differentiation phase, the cells were incubated for another 5 days in PGM containing G-CSF only. To trigger differentiation towards monocytes, the cells were first grown for 3 days in PGM containing IL3+SCF. They were then incubated for 2 days in the presence of M-CSF (macrophage colony stimulating factor, a.k.a. CSF1) and GM-CSF (granulocyte/macrophage colony stimulating factor, a.k.a. CSF2), and finally cultivated for 5 days in the presence of M-CSF only. Cell density was maintained at $2x10^5$ cells/mL throughout the first 5 days of culture, both for neutrophil and monocyte differentiation. All cytokines were from PeproTech (Hamburg, Germany) and were used at a final concentration of 50ng/mL.
To study the effect of iron complementation on the differentiation of P1/2-KO progenitor cells, ferric ammonium citrate (Sigma-Aldrich) was added to the culture medium during the G-CSF+IL3 and G-CSF alone

treatment periods. To study the consequences of iron deprivation on P1/2-CTR cell differentiation, deferiprone (Sigma-Aldrich) was added to the culture medium also during the G-CSF+IL3 and G-CSF exposure phases. Control cells not subjected to iron treatments were exposed to vehicle.

### QUANTIFICATION AND STATISTICAL ANALYSES

#### Transcriptome analysis

Protein coding genes with a $\log_2$(fold change) > 1 or <-1 (FDR 10%) in any of the pairwise comparisons tested (PMN versus Pro, PrNeu versus Pro, PMN versus PrNeu in P1/2-CTR samples and separately in P1/2-KO) were selected for subsequent analysis. For each gene, TPM (transcripts per million) values across the three cell populations tested were scaled to generate z-scores. The z-scores were then used to define specific expression patterns across the Pro, prNeu and PMN differentiation stages (i.e. during differentiation), and the genes were grouped in 4 distinct clusters using the k-means method with the Hartigan-Wong algorithm (Hartigan and Won, 1979). Genes whose expression in P1/2-KO cells did not follow the same pattern as in P1/2-CTR were subsequently analyzed for GO (gene ontology)-term enrichment using the topGO package (Alexa and Rahnenfuhrer, DOI: 10.18129/B9.bioc.topGO), and were plotted using the ggplot2 package (bioconductor). Heatmaps were generated on scaled expression values using the heatmap.2 function of the ggplot2 package.

#### Proteome analysis

The statistical analysis was performed with the R-package “limma” (Ritchie et al., 2015). The LFQ (Cox etal., 2014) values were used for the statistical analysis. Adapted from the Perseus recommendations (Tyanova et al., 2016b) protein groups with non-zero intensity values in 70% of the samples of at least one of the conditions were used and for missing values being completely absent in one condition imputation with random values drawn from a downshifted (2.2 standard deviation) and narrowed (0.3 standard deviation) intensity distribution of the individual sample was applied. For missing values with no complete absence in one condition the R package missForest was used for imputation (Stekhovenand Buehlmann, 2012). The p-values were adjusted with the Benjamini–Hochberg method for multiple testing (Benjamini and Hochberg, 1995).

Perseus (version 1.6.15.0) was used for mouse protein annotation with gene ontology (GO) and KEGG and 1D enrichment analysis (Cox and Mann, 2012). The 1D enrichment was performed with a Benjamini-Hochberg FDR of 0.02. The 1D enrichment on log fold changes accepting changes to both sides and the 1D enrichment on adjusted p-values accepting only changes to lower.

#### Other data

The values are presented as box plots with minimum to maximum values, or as mean ± standard error of the mean, as indicated in Figure legends. The normality of two independent groups of data was first assessed using the D'Agostino-Pearson test and then compared using either a two-tailed unpaired Student t test or a Mann-Whitney test. In all statistical analyses, the null hypothesis was rejected for p-values below 0.05. All analyses were performed using the Prism application (v9) from GraphPad (GraphPad Software, San Diego, CA).

Figure S1

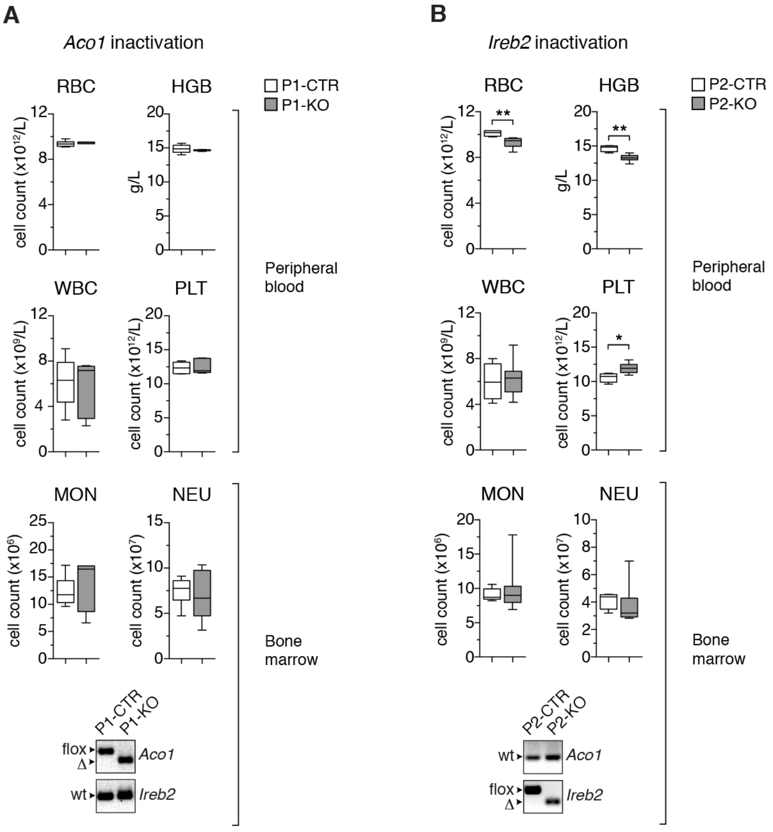


Figure S1: **Acute ablation of either IRP1 or IRP2 does not impair steady-state myelopoiesis**, related to Figure 1.

(A) Adult *Aco1*$^{flox/flox}$*Rosa26*$^{+/CreERT2}$ (designated P1-KO) and *Aco1*$^{flox/flox}$*Rosa26*$^{+/+}$ (P1-CTR) mice (n=5-6) were treated with tamoxifen on day 1 and day 3 to induce IRP1 ablation. They were sacrificed on day 10 to analyze hematological PB parameters (RBC: red blood cell; WBC: white blood cells; PLT: platelets, HGB: hemoglobin), and the amount of monocytes (MON) and neutrophils (NEU) in the bone marrow (BM) using flow cytometry (FCM, cell counts in the two hindlimbs).

(B) Adult *Ireb2*$^{flox/flox}$*Rosa26*$^{+/CreERT2}$ (P2-KO) and *Ireb2*$^{flox/flox}$*Rosa26*$^{+/+}$ (P2-CTR) mice (n=4-7) were subjected to the same treatment to assess the effect of acute ablation of IRP2.

Acute IRP1 disruption had no significant effect on RBC counts during adulthood. Acute IRP2 ablation caused mild anemia, associated with a slight increase in PLT counts, which was reminiscent of the anemia observed in mice with constitutive deletion of IRP2 (Cooperman et al., 2005, Galy et al., 2005b). In sharp contrast to the myeloid phenotype of P1/2-KO mice lacking both IRP1 and IRP2, neither IRP1 deficiency nor IRP2 disruption affected WBC or neutrophil counts in the BM. Genomic PCR analysis confirmed efficient recombination of Irp1 (*Aco1*) and Irp2 (*Ireb2*) floxed alleles in BM cells of P1-KO and P2-KO mice, respectively (lower panels; flox: floxed allele; Δ: truncated allele). The results are presented as box plots with minimum to maximum values. * $p<0.05$; ** $p<0.01$.

*See also Figure 1.*

Figure S2

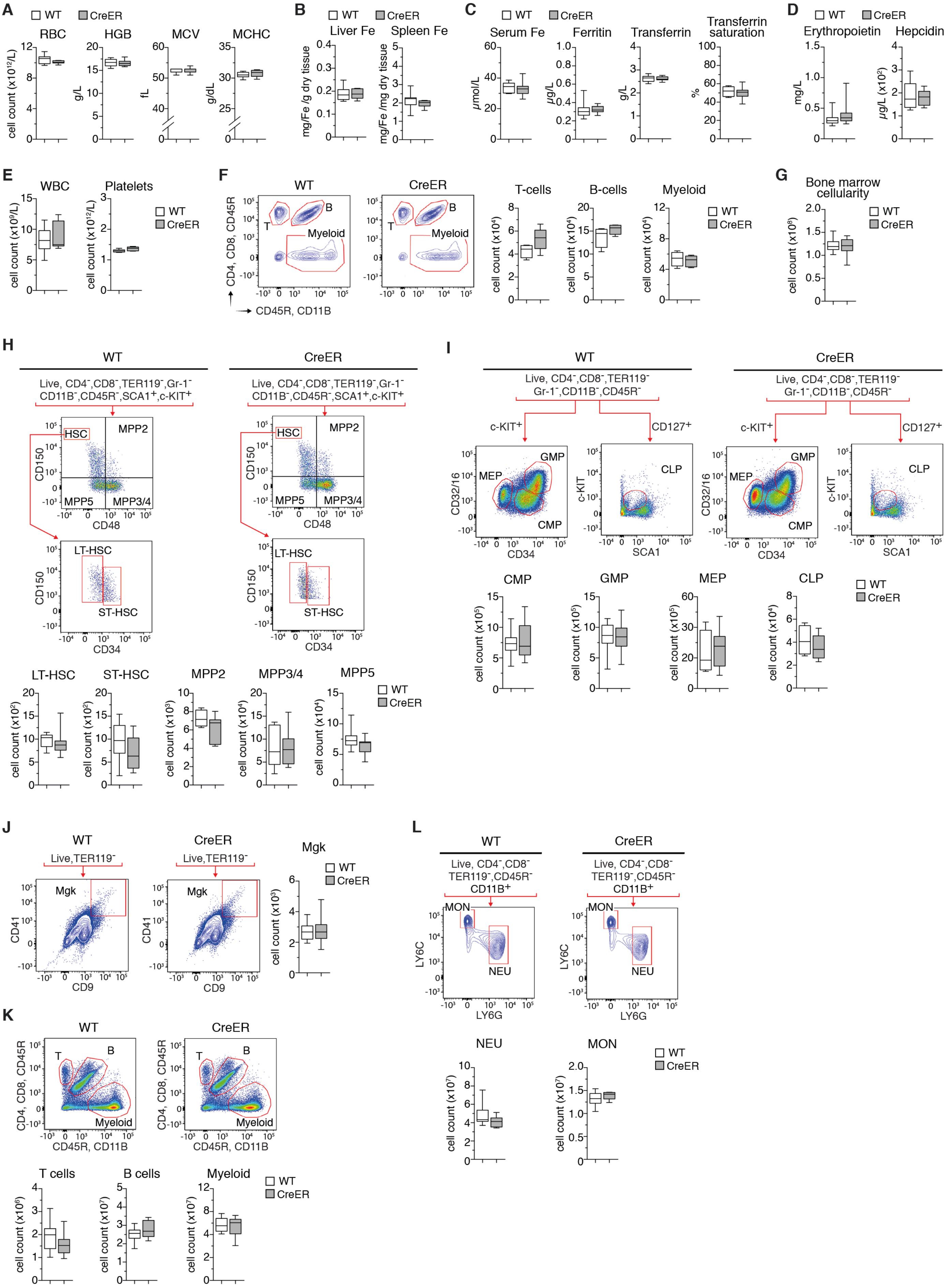
A
WT
CreER
RBC
HGB
MCV
MCHC
cell count (x10^12/L)
g/L
fL
g/dL
B
Liver Fe
Spleen Fe
mg/Fe /g dry tissue
mg/Fe /mg dry tissue
C
Serum Fe
Ferritin
Transferrin
Transferrin saturation
µmol/L
µg/L
g/L
%
D
Erythropoietin
Hepcidin
mg/L
µg/L (x10^2)
E
WBC
Platelets
cell count (x10^9/L)
cell count (x10^12/L)
F
T
B
Myeloid
CD4, CD8, CD45R
CD45R, CD11B
T-cells
B-cells
cell count (x10^4)
G
Bone marrow cellularity
cell count (x10^8)
H
Live, CD4-,CD8-,TER119-,Gr-1-
CD11B-,CD45R-,SCA1+,c-KIT+
HSC
MPP2
MPP5
MPP3/4
CD150
CD48
LT-HSC
ST-HSC
CD34
cell count (x10^2)
cell count (x10^3)
I
Live, CD4-,CD8-,TER119-
Gr-1-,CD11B-,CD45R-
c-KIT+
CD127+
GMP
MEP
CMP
CLP
CD32/16
c-KIT
SCA1
cell count (x10^5)
J
Live,TER119-
Mgk
CD41
CD9
K
T cells
B cells
cell count (x10^6)
cell count (x10^7)
L
Live, CD4-,CD8-
TER119-,CD45R-
CD11B+
MON
NEU
LY6C
LY6G

Figure S2: **Effect of CRE activation on blood cell counts, serum iron parameters, and bone marrow cell populations**, related to Figures 1 and 2.
WT and CreER littermates received tamoxifen on day 1 and day 3 to activate CRE. Mice were analyzed on day 10.
(A) Red blood cell (RBC) parameters (HGB: hemoglobin; MCV: mean corpuscular volume; MCHC: mean corpuscular hemoglobin concentration) (n=7-11).
(B) Hepatic and splenic iron levels (n=7-11).
(C) Serum iron parameters (n=7-11).
(D) Serum levels of erythropoietin and hepcidin (n=7-11).
(E) White blood cell (WBC) and platelet counts in peripheral blood (PB) (n=7-11).
(F) Flow cytometry (FCM) analysis of major WBC populations in PB. The gating strategy is shown on contour plots on the left. The histograms display cell counts for $3x10^5$ events recorded (n=4-5)
(G) Bone marrow (BM) cellularity in WT versus CreER littermates (n=10-14).
H-L) FCM analysis of BM cell populations including: (H) long term (LT)- and short term (ST)-HSCs (hematopoietic stem cells), multipotent progenitors (MPP, n=6-13), (I) common myeloid (CMP) and lymphoid (CLP) progenitors, megakaryocyte/erythroid progenitors (MEP), and granulocyte/macrophage (GMP) progenitors (n=6-14); (J) megakaryocytes (Mgk) (n=6-8); (K) T- and B-lymphocytes, and myeloid cells (granulocytes/monocytes) (n=10-14); (L) monocytes versus neutrophils (n=7-8). The gating strategy is indicated for each cell population analyzed. Box plots (minimum to maximum values) show the number of cells present in the BM of both hindlimbs.
* p<0.05; ** p<0.01; *** p<0.001). CRE activation in CreER mice does not have any obvious effect on the parameters measured .
*See also Figures 1 and 2.*

Figure S3

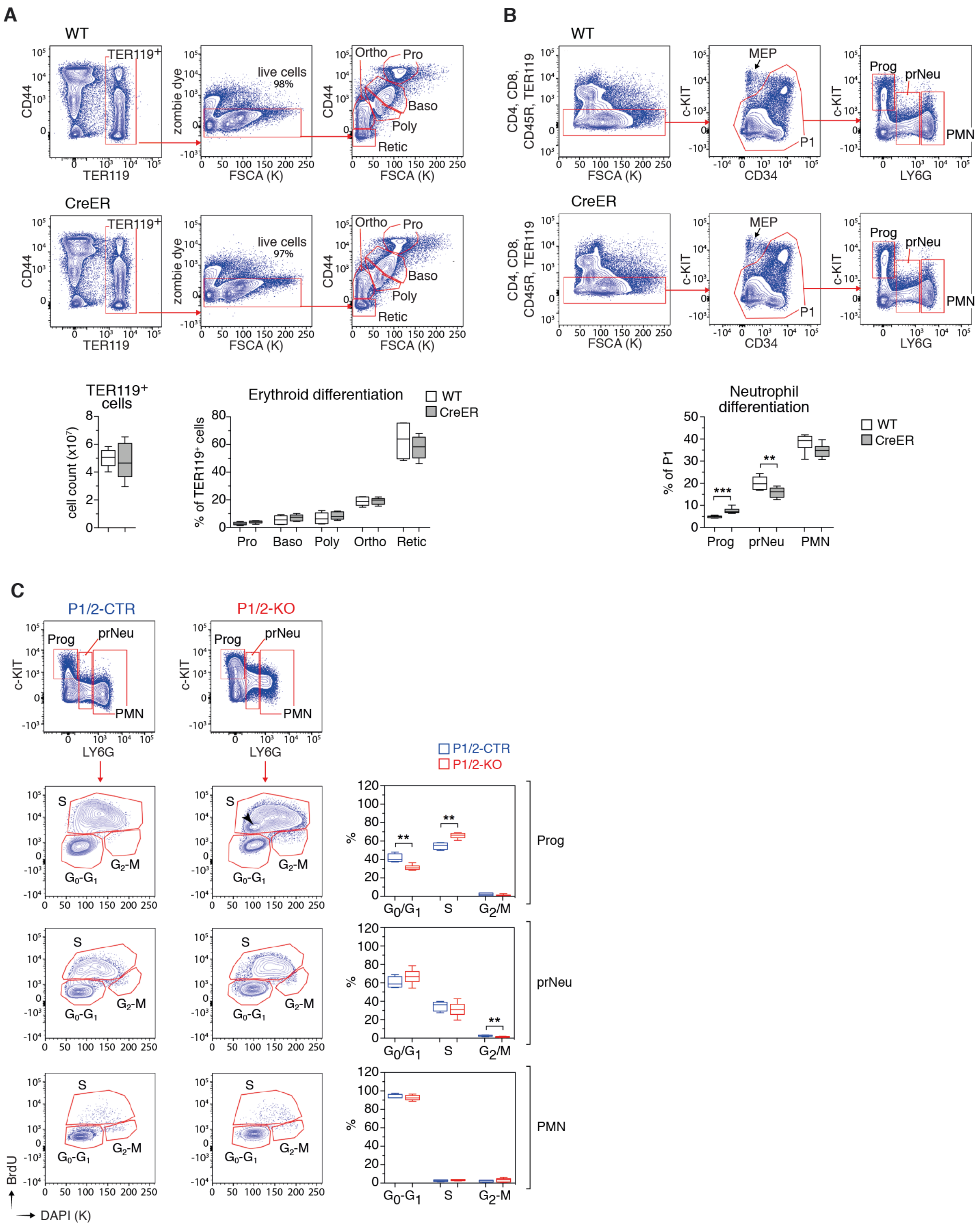


Figure S3: **Effect of CRE on erythroid and neutrophil differentiation in the bone marrow, and consequences of IRP deficiency on the cell cycle during neutropoiesis**, related to Figure 3.

(A) FCM analysis of terminal erythroid differentiation in the BM of CreER versus WT mice. Five TER119$^+$ cell populations corresponding to pro-erythroblasts (Pro), basophilic (Baso), poly- (Poly) and orthro-chromatic (Ortho) cells, and reticulocytes (Retic) are gated based on cell size (FSCA: forward scatter area) and the CD44 marker. Box plots (minimum to maximum values, n=5-6) show the number of TER119$^+$ cells present in the

BM of both hindlimbs (left), and the frequency of cells at different stages of differentiation (right). CRE activation in CreER mice did not alter the number of TER119+ cells or terminal differentiation of erythrocytes.
(B) FCM analysis of neutrophil differentiation in the BM of CreER versus WT mice; c-KIT$^{high}$LY6G$^{-}$ cells represent progenitor (Prog) cells; c-KIT$^{-}$LY6G$^{low}$, pre-neutrophil stage (prNeu); c-KIT$^{-}$LY6G$^{high}$, polymorphonuclear (PMN) neutrophils. The box plot (minimum to maximum values, n=7) displays the overall frequency of Prog, prNeu, and PMN cells in the BM. Activation of CRE triggered a minimal, 1.5-fold increase in the amount of Prog cells in the BM, which was significantly less than the 4.2-fold increase observed in P1/2-KO mice (see Figure 3B). CreER mice also display a 20% decrease in prNeu cells that is not observed in P1/2-KO mice, and had normal PMN counts.
(C) FCM analysis of cell cycle progression during neutropiesis in P1/2-KO versus P1/2-CTR mice, based on *in vivo* labeling of DNA with the nucleoside analogue 5-bromo,2-deoxyuridine (BrdU). Representative FCM plots are shown (left). The top FCM panels show the gating of Prog, prNeu and PMN cells based on LY6G and c-KIT staining. The lower panels display the gating of cells in the $G_0/G_1$, S, and $G_2$/M phases of the cell cycle, respectively. Box plots on the right (minimum to maximum values) show the percentage of cells in the different phases of the cell cycle (P1/2-KO, n=11, P1/2-CTR mice n =4). As expected, the frequency of proliferating cells decreased from the Prog to the PMN stage, with nearly all PMN cells in $G_0/G_1$. The most noticeable difference between P1/2-KO and P1/2-CTR mice was an increase in the proportion of Prog cells in the S phase, associated with a reduction of cells in $G_0/G_1$ (top). A small proportion of P1/2-KO Prog cells incorporated BrdU but seem stopped in the S phase (black arrowhead).
(A-D) the p-value corresponds to separate pairwise comparisons between P1/2-CTR and P1/2-KO for each parameter and cell population analyzed, ** p<0.01; *** p<0.001.
*See also Figure 3.*

Figure S4

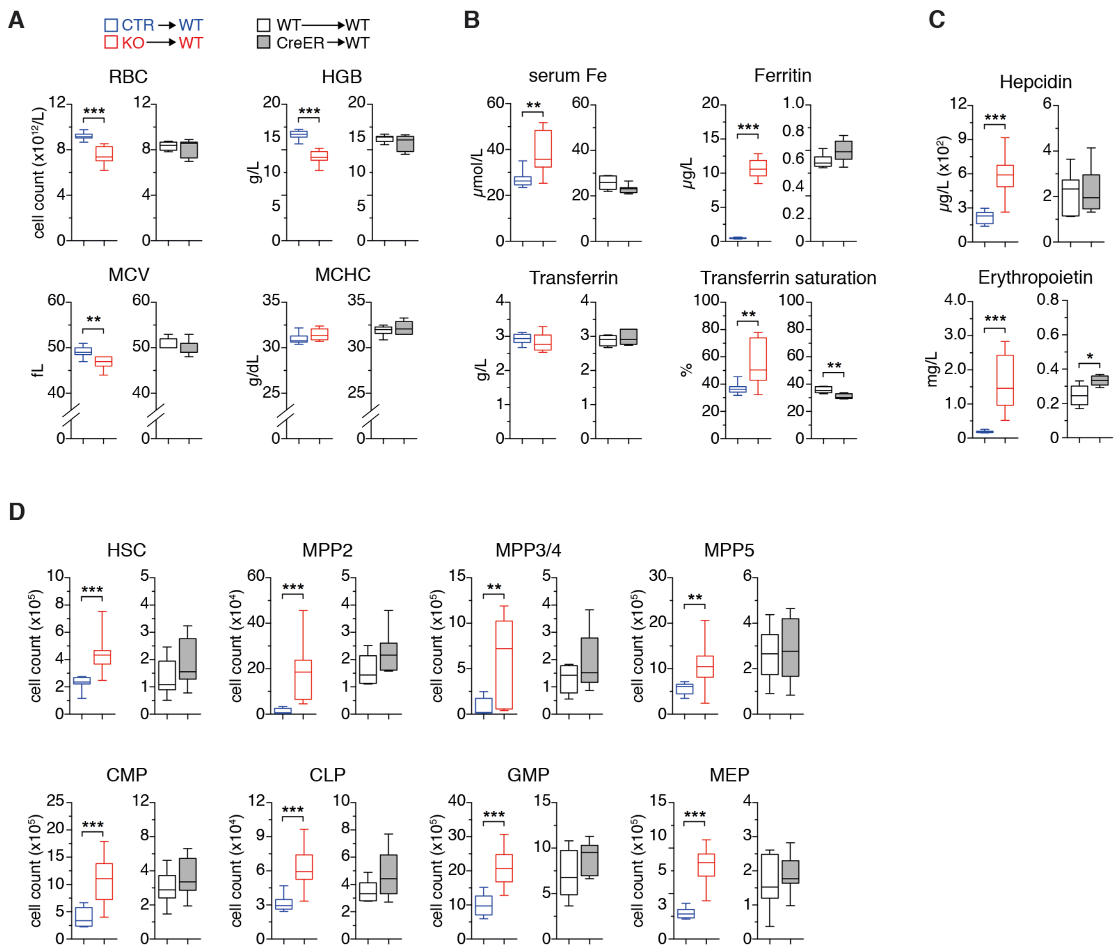


Figure S4: **Mice lacking IRP expression in hematopoietic cells phenocopy mice with systemic loss of IRP function**, related to Figure 4.

As in Figure 4, BM cells from P1/2-KO mice were transplanted into WT recipients to inactivate the IRP/IRE system in hematopoietic cells only (chimeras designated KO→WT). As a control, recipients were transplanted with BM cells from P1/2-CTR animals (CTR→WT). To identify possible effects of CRE, recipients were also transplanted with BM cells from CreER (CreER→WT) or WT donors (WT→WT). Following stable engraftment, the chimeras were treated with tamoxifen on day 1 and day 3, and were sacrificed on day 10 for the analysis of (A) red blood cell (RBC) indices in peripheral blood (HGB: hemoglobin; MCV: mean corpuscular volume; MCHC: mean corpuscular hemoglobin concentration), (B) plasma iron parameters, (C) hepcidin and erythropoietin (EPO) levels in the circulation, and (D) the amount of $LIN^-$ cell populations in the BM of both hindlimbs.

Loss of IRP function in hematopoietic cells of KO→WT chimeras caused microcytic anemia (A), associated with high plasma levels of iron and ferritin (B) and high hepcidin and EPO values (C). KO→WT mice also displayed an expansion of all $LIN^-$ cell compartments in the BM. Those features recapitulate the main phenotypic alterations observed in P1/2-KO mice with acute, systemic ablation of IRP1 and IRP2 (compare with Figures 1 and 2). Of note, activation of CRE alone in hematopoietic cells of CreER→WT chimeras did not cause the microcytic anemia, hyperferremia, and expansion of $LIN^-$ cells seen in P1/2-KO and KO→WT animals. The results are displayed as box plots with minimum to maximum values (n=7-10). ** $p<0.01$; *** $p<0.001$.

*See also Figure 4.*

Figure S5

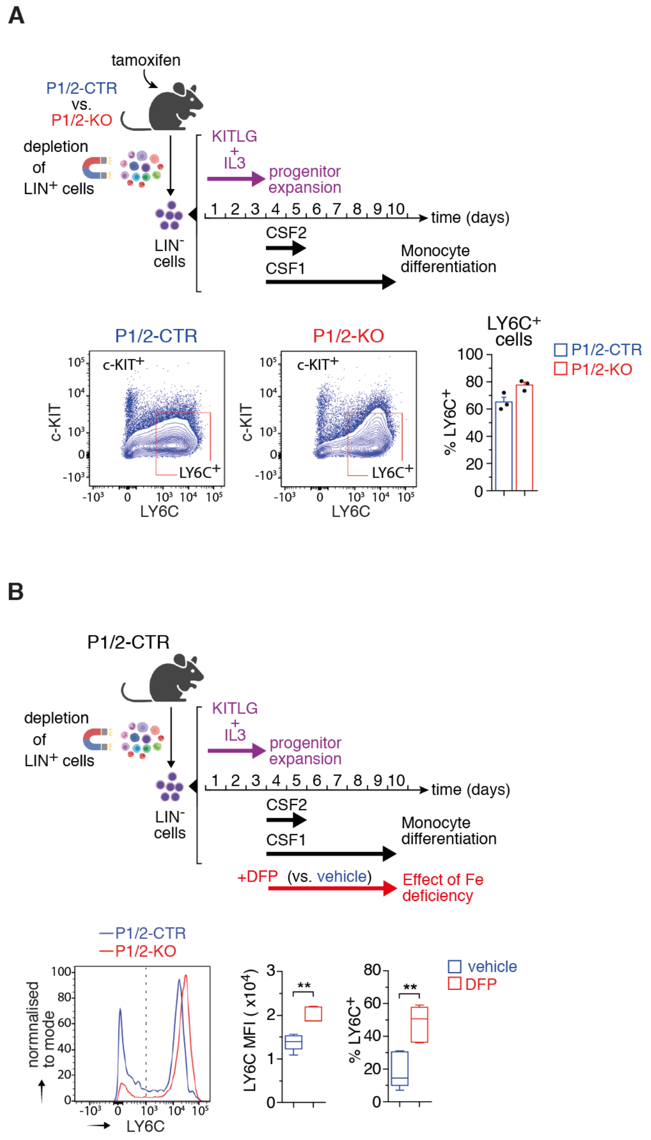


Figure S5: **Effects of IRP deficiency and iron deprivation on the differentiation of hematopoietic progenitor cells towards monocytes**, related to Figures 4 and 7.

(A) Top: Lineage-negative (LIN$^-$) cells from the BM of tamoxifen-treated P1/2-CTR and P1/2-KO mice were expanded *ex vivo* in the presence of KIT ligand (KITLG) and interleukin 3 (IL3). They were subsequently differentiated into LY6C$^+$ monocytes first with CSF2+CSF1 and then with CSF1 alone, as indicated. Bottom: representative FCM plots showing LY6C versus c-KIT levels and the gating of LY6C$^+$ cells; the bar graph (mean +SEM) shows the % of LY6C$^+$ cells obtained with BM progenitors from 3 animals for each genotype. IRP deficiency did not impair the differentiation of hematopoietic progenitor cells towards monocytes, in sharp contrast to its effect on differentiation towards neutrophils (see figure 4D).

(B) Top: LIN$^-$ cells from the BM of untreated P1/2-CTR control mice were differentiated towards monocytes as described above but in the presence of 50µM of the iron chelator deferiprone (DFP) versus vehicle. Bottom: Representative FCM plot showing LY6C staining; the dotted line delimits LY6C$^+$ from LY6C$^-$ cells. Box plots (minimum to maximum values, n=5) display the median fluorescence intensity (MFI) and percentage of LY6C$^+$ cells. Opposite to its inhibitory effect on LY6G$^+$ neutrophils (see Figure 4D), the iron chelator enhanced LY6C expression and the differentiation of BM progenitor cells towards LY6C monocytes. ** $p<0.01$).

(A-B) top schemes created with BioRender.com.

*See also Figures 4 and 7.*

Figure S6

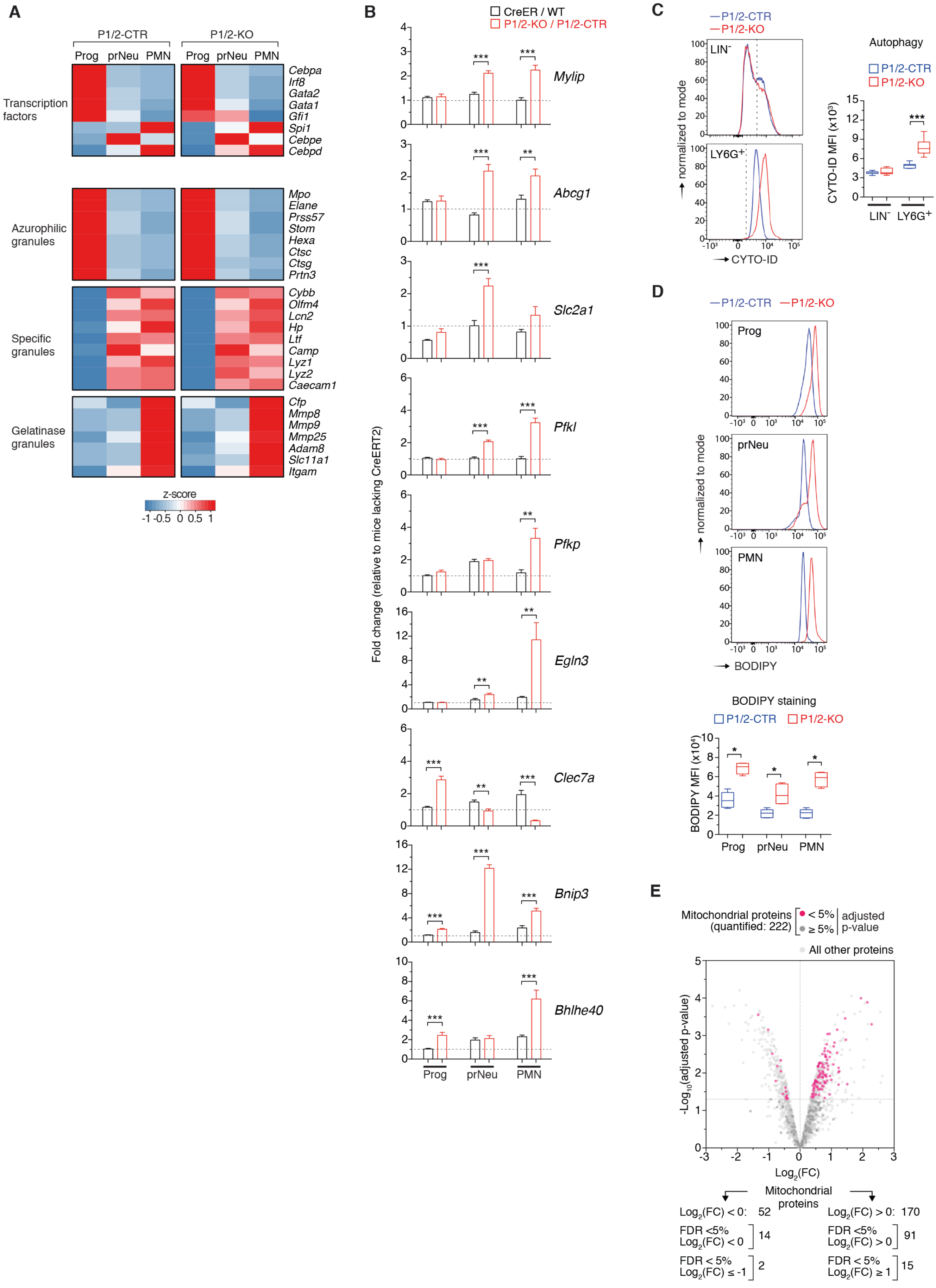
A
P1/2-CTR
P1/2-KO
Prog prNeu PMN
Prog prNeu PMN
Transcription factors
Cebpa
Irf8
Gata2
Gata1
Gfi1
Spi1
Cebpe
Cebpd
Azurophilic granules
Mpo
Elane
Prss57
Stom
Hexa
Ctsc
Ctsg
Prtn3
Specific granules
Cybb
Olfm4
Lcn2
Hp
Ltf
Camp
Lyz1
Lyz2
Caecam1
Gelatinase granules
Cfp
Mmp8
Mmp9
Mmp25
Adam8
Slc11a1
Itgam
z-score
-1 -0.5 0 0.5 1
B
CreER / WT
P1/2-KO / P1/2-CTR
Fold change (relative to mice lacking CreERT2)
Mylip
Abcg1
Slc2a1
Pfkl
Pfkp
Egln3
Clec7a
Bnip3
Bhlhe40
Prog prNeu PMN
C
P1/2-CTR
P1/2-KO
LIN-
LY6G+
normalized to mode
CYTO-ID
Autophagy
P1/2-CTR
P1/2-KO
CYTO-ID MFI (x10³)
LIN- LY6G+
D
P1/2-CTR
P1/2-KO
Prog
prNeu
PMN
normalized to mode
BODIPY
BODIPY staining
P1/2-CTR
P1/2-KO
BODIPY MFI (x10⁴)
Prog prNeu PMN
E
Mitochondrial proteins (quantified: 222)
< 5%
≥ 5%
adjusted p-value
All other proteins
-Log10(adjusted p-value)
Log2(FC)
Mitochondrial proteins
Log2(FC) < 0: 52
Log2(FC) > 0: 170
FDR <5% Log2(FC) < 0 14
FDR <5% Log2(FC) > 0 91
FDR < 5% Log2(FC) ≤ -1 2
FDR < 5% Log2(FC) ≥ 1 15

Figure S6: **Validation of the analysis and interpretation of transcriptome and proteome data**, related to Figure 5.

(A) Heat maps representing the expression trajectory of selected mRNAs during neutropoiesis in P1/2-CTR and P1/2-KO cells (based on RNA-seq data). The mRNAs selected encode transcription factors important for myelopoiesis (top) as well as proteins associated with azurophylic, specific, and gelatinase granule formation. The expression trajectory of these genes is very similar in P1/2-CTR and P1/2-KO cells.

(B) qRT-PCR analysis of selected mRNAs in Prog, prNeu and PMN cells isolated from the BM of P1/2-KO versus P1/2-CTR mice. The bar graphs show the expression ratio (mean ±SEM, n=3) in P1/2-KO versus P1/2-CTR samples (red bars) after calibration on the average expression of four standard transcripts (*Gusb*, *Ppib*, *Ubxn4*, and *Psmc2*). Several mRNAs were expressed at an abnormally high level in LY6G$^+$ P1/2-KO cells compared with P1/2-CTR, confirming the RNA-seq analysis (see Figure 5). The mRNAs selected for the analysis are involved in glycolysis (*Slc2a1*, *Pfkl*, *Pfkp*), the regulation of transcription (*Bhlhe40*), oxygen sensing (*Egln3*), cell survival (Bnip3), or cholesterol metabolism (*Abcg1*, *Mylip*). Also consistent with the RNA-seq data, *Clec7a* (pattern-recognition receptor), was overexpressed in Prog cells but was downregulated in PMN cells. Cells from CreER versus WT animals were analyzed in parallel (black bars). Although CRE activation had a slight effect on some transcripts (e.g. slightly elevated *Pfkp* levels in PrNeu cells), most expression changes observed are clearly attributed to IRP ablation.

(C) FCM analysis with CYTO-ID dye was used to analyze autophagic vacuoles in whole BM LIN$^-$ and LY6G$^+$ cells from P1/2-KO versus P1/2-CTR mice. Top: representative FCM plot. Bottom: box plot (minimum to maximum values, n=7) displaying CYTO-ID MFI. LY6G$^+$ cells from P1/2-KO mice exhibited significantly higher activity of the autophagic pathway compared to control; IRP ablation did not have a detectable impact on autophagic activity in LIN$^-$ cells.

(D) FCM analysis of lipid droplets during neutrophil differentiation with the BODIPY 493/503 dye. Representative FCM plots are shown together with a box plot (minimum to maximum values, n=4) displaying BODIPY MFI. This analysis revealed an elevation of neutral lipid levels in Prog, prNeu and PMN cells from P1/2-KO mice.

(E) Volcano plot showing the expression of proteins in whole BM LY6G$^+$ positive cells from P1/2-KO versus P1/2-CTR mice (all proteins quantified are represented). FC (fold change): P1/2-KO – P1/2-CTR. The 222 proteins highlighted are those listed in the Mitocarta (v3.0) database (Rath et al., 2021). The color code indicates the p-value adjusted with the Benjamini–Hochberg method for multiple testing. The majority of mitochondrial proteins tend to be expressed at a higher level in P1/2-KO cells.

(C,D): The dotted line delineates marker-positive and -negative cells.

(B-D): p-values correspond to pair-wise comparisons between P1/2-CTR and P1/2-KO for each parameter/cell population tested. * $p<0.05$; ** $p<0.01$; *** $p<0.001$.

*See also Figure 5.*

Figure S7

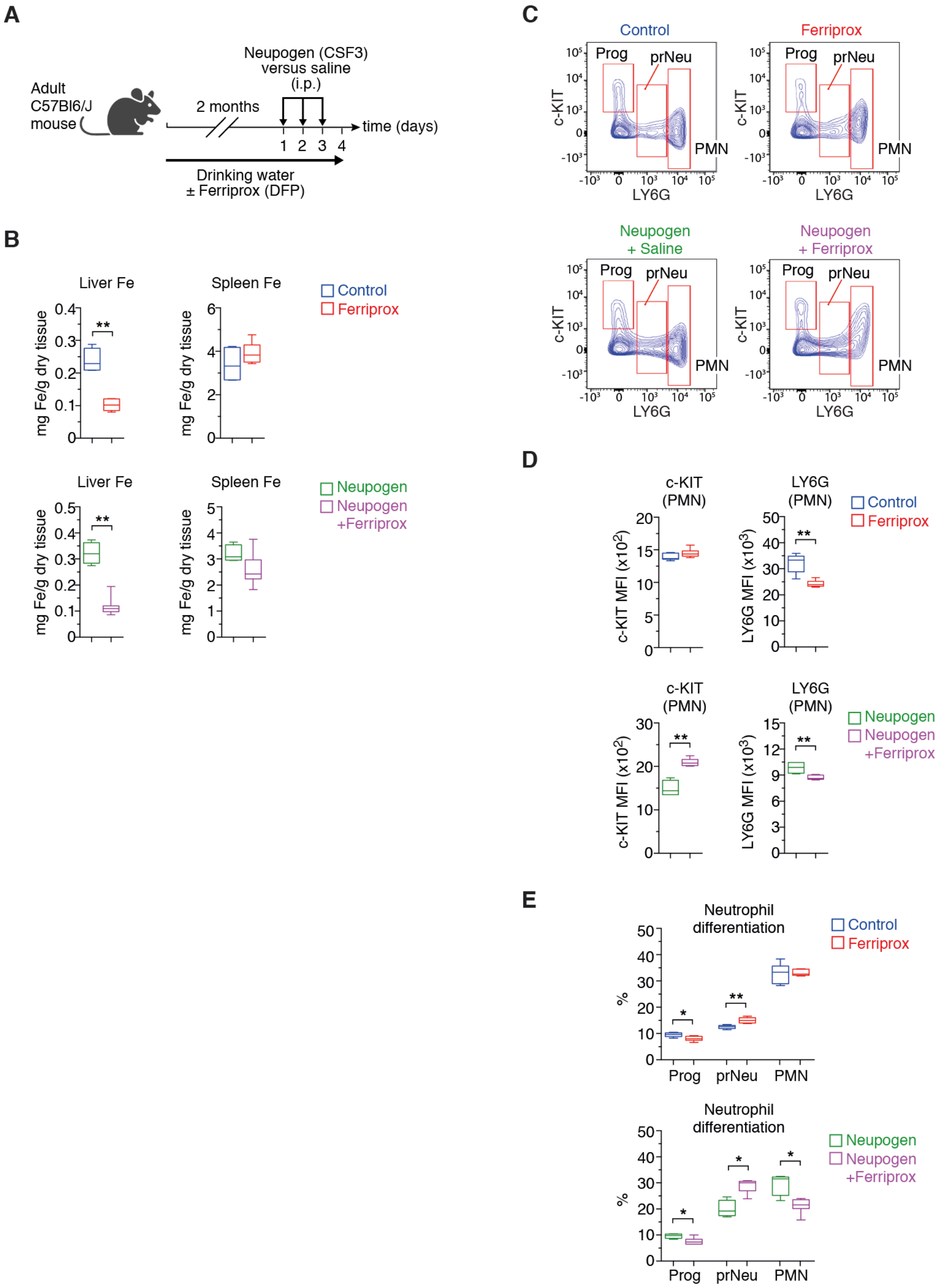


Figure S7: **Effect of chronic treatment with the iron chelator deferiprone on steady-state and emergency neutropoiesis *in vivo***, related to Figure 7.

(A) Wild type adult mice were given deferiprone (DFP) in the form of Ferriprox (added to the drinking water) for 2 months; control mice were given regular drinking water. At the end of the DFP treatment, they were injected (i.p.) daily with CSF3 (G-CSF, in the form of Neupogen) on 3 consecutive days to stimulate granulopoiesis and were analyzed one day after the last CSF3 injection; control animals received saline only. Created with BioRender.com.

(B) Effect of DFP/Ferriprox on hepatic and splenic iron levels under baseline conditions (top) or upon stimulation of granulopoiesis with CSF3/Neupogen (bottom).

(C) Representative FCM plots showing c-KIT and LY6G levels in Prog, prNeu, and PMN cells in the BM.

(D) Box plots (minimum to maximum values) displaying the median fluorescence intensity (MFI) of the c-KIT and LY6G markers in PMN cells.
(E) Box plot showing the percentage of Prog, PrNeu and PMN cells in the BM; stimulation of granulopoiesis with CSF3/Neupogen increased the proportion of prNeu cells in mice receiving either regular water or DFP/Ferriprox.
DFP/Ferriprox alone had a minor effect on basal neutropoiesis, with a slight but significant decrease in LY6G MFI in PMN cells (D, top). In mice stimulated with CSF3/Neupogen, iron chelation with DFP/Ferriprox led to a reduction of the proportion of PMN cells (E, bottom). Those PMN cells were characterized by higher expression of c-KIT and lower levels of LY6G (D, bottom); reminiscent of the c-KIT$^{+}$LY6G$^{+}$ PMN cell population observed in the BM of P1/2-KO mice (cf. Figure 3B). Chronic iron chelation with DFP/ Ferriprox had marginal effects on basal neutropoiesis, however, significantly altered neutrophil differentiation in the BM under conditions of emergency granulopoiesis.
(B,D,E) For each parameter and cell population analyzed, p-values correspond to separate pairwise comparisons between the DFP/Ferriprox group (n=6) and the control group (n=6), or between the CSF3/Neupogen+saline (n=4) and the CSF3/Neupogen+DFP/Ferriprox (n=7) groups. * p<0.01; ** p<0.01.
*See also Figure 7.*
